# Characterization of a Novel Cell Wall-Associated Nucleotidase of *Enterococcus faecalis* that Degrades Extracellular c-di-AMP

**DOI:** 10.1101/2025.06.08.658492

**Authors:** Adriana G Morales Rivera, Anju Bala, Leila G Casella, Debra N Brunson, Aria Patel, Elsa Wongso, Ana L Flores-Mireles, José A Lemos

## Abstract

*Enterococcus faecalis* is a prolific opportunistic pathogen responsible for a range of life-threatening infections for which treatment options are increasingly limited due to the high prevalence of multidrug-resistant isolates. Cyclic di-AMP has emerged as an essential bacterial signaling molecule due to its impact on physiological processes, including osmotic adaptation, cell wall homeostasis, antibiotic tolerance, and virulence. In addition, c-di-AMP is a potent pathogen-associated molecular pattern (PAMP) molecule recognized by the host immune system to trigger protective responses. In previous work, we identified and characterized the enzymes responsible for the synthesis and degradation of intracellular c-di-AMP in *E. faecalis*, demonstrating that maintaining c-di-AMP homeostasis is vital for bacterial fitness and virulence. In addition to the intracellular enzymes that regulate c-di-AMP levels, a limited number of bacteria encode surface-associated nucleotidases capable of cleaving extracellular c-di-AMP, potentially facilitating immune evasion. Here, we characterize a novel and unique cell wall-anchored phosphodiesterase, termed EecP (*E. faecalis* extracellular c-di-AMP phosphodiesterase), which features duplicated catalytic domains and specifically degrades extracellular c-di-AMP. Deletion of *eecP* (Δ*eecP*) resulted in a marked accumulation of extracellular c-di-AMP. Although the Δ*eecP* strain exhibited comparable growth and behavior to the parent strain *in vitro*, it displayed increased susceptibility to killing by phagocytic cells. Using two murine infection models, we show that the impact of *eecP* deletion and the consequent buildup of extracellular c-di-AMP on *E. faecalis* pathogenesis may be site-specific. Notably, disseminated infection was more severe in mice infected with the Δ*eecP* strain, suggesting that extracellular c-di-AMP influences infection outcomes, likely through modulation of host immune responses.

**Author Summary:** *Enterococcus faecalis* is a major opportunistic pathogen and a leading cause of several life-threatening hospital-associated infections. Cyclic di-AMP is a bacterial second messenger nucleotide that regulates essential cellular processes and plays key roles in bacterial pathogenesis and host immune activation. We previously characterized the enzymes responsible for the synthesis and degradation of c-di-AMP in *E. faecalis*, demonstrating that this signaling molecule is crucial for bacterial fitness and virulence. In this study, we describe the characterization of EecP, a novel cell wall-associated enzyme that degrades c-di-AMP extracellularly. Our findings identify EecP as a new virulence factor in *E. faecalis*, capable of modulating infection outcomes.

## Introduction

*Enterococcus faecalis* is a Gram-positive facultative anaerobe and commensal inhabitant of the gastrointestinal tract of humans and animals. Notorious for its prevalence as a major healthcare-associated opportunistic pathogen, *E. faecalis* was responsible for an estimated toll of over 200,000 worldwide deaths in 2019 alone^1^. Although generally considered a low-grade pathogen due to its limited repertoire of tissue-damaging virulence factors, the pathogenic potential of *E. faecalis* largely stems from its ability to form robust biofilms on medical devices, evade the immune system, and survive under different stressful conditions. These factors collectively enable its persistence in hospital environments and contribute to opportunistic infections such as catheter-associated urinary tract infections (CAUTI), intra-abdominal and soft tissue infections, and central line-associated bloodstream infections^2,3^. Moreover, *E. faecalis* possesses intrinsic and acquired resistance to multiple antibiotics, drastically complicating treatment and worsening prognosis^4^.

Since its serendipitous discovery in the crystal structure of the DNA integrity scanning protein DisA from *Thermotoga maritima* in 2008^5^, bis-(3’-5’)- cyclic dimeric adenosine monophosphate (c-di-AMP) has emerged as an essential nucleotide second messenger for bacteria required for the regulation of numerous cellular functions, including osmotic adaptation, DNA repair, cell wall homeostasis, antibiotic tolerance, and virulence^6–12^. Cyclic di-AMP is synthesized from two adenosine triphosphate (ATP) molecules by diadenylate cyclase (DAC) enzymes and degraded to phosphoadenylyl adenosine (5’ pApA) or adenosine monophosphate (AMP) by enzymes known as phosphodiesterases (PDE). As a ligand, c-di-AMP can bind allosterically to effector proteins and positively or negatively modulate their activity^6–12^. To date, over twenty targets have been identified to be under c-di-AMP allosteric control, including membrane-associated transporters, signal transduction proteins, metabolic enzymes, regulatory proteins, and riboswitches^6–12^.

While essential for cellular functions, bacteria must tightly control intracellular c-di-AMP pools by modulating the activities of DAC and PDE enzymes and, arguably, through active secretion mechanisms^13^. Bacterial strains that cannot synthesize c-di-AMP are frequently only viable under particular conditions, with several examples indicating that c-di-AMP production is essential for bacterial virulence^14–16^. At the same time, PDE mutants that accumulate intracellular c-di-AMP also display important phenotypes, from altered stress responses to attenuated virulence in mouse models ^17–19^. Previously, our group characterized the enzymes that synthesize (CdaA) and degrade (DhhP and GdpP) c-di-AMP in *E. faecalis*^15^. The key finding of this study was that disruption of c-di-AMP homeostasis, either the intracellular accumulation observed in strains lacking one or two PDEs (*gdpP* and *dhhP*) or complete lack of c-di-AMP observed in the Δ*cdaA* strain, drastically impaired *E. faecalis* fitness and virulence^15^.

Beyond its role in bacterial physiology, c-di-AMP is also a major pathogen-associated molecular pattern (PAMP) molecule recognized by host immune and tissue cells^9, 20–22^. Specifically, c-di-AMP is recognized by at least four host cell receptors, namely STING (<u>st</u>imulator of <u>in</u>terferon <u>g</u>enes)^20, 21, 23^, DDX41 (DEAD-box RNA helicase 41)^24^, RECON (reductase controlling NF-κB or AKR1C13)^25^, and ERAdP (endoplasmic reticulum membrane adaptor)^26^. These receptors activate various innate immune signaling pathways and promote the production of cytokines that modulate the immune response against bacteria. As such, c-di-AMP sensing is thought to play a key role in antibacterial host responses and limiting infection progression.

In addition to cytoplasmic PDEs, a few bacteria have been shown to encode surface-associated enzymes, termed extracellular phosphodiesterases (ePDE), that hydrolyze extracellular c-di-AMP. The first description of an ePDE was in the human pathogen *Streptococcus agalactiae* with the identification of the cell wall-anchored enzyme CdnP^20^. In this study, CdnP was shown to degrade extracellular c-di-AMP, dampening activation of the STING-mediated response, leading the authors to propose that CdnP serves as an immune evasion strategy for *S. agalactiae*^20^. A second c-di-AMP ePDE, SntA, initially characterized as a cell wall-anchored heme-binding protein that interferes with complement-mediated killing, was later identified in the zoonotic pathogen *Streptococcus suis*^27, 28^. Biochemical studies confirmed that SntA can efficiently degrade c-di-AMP as well as other nucleotides, including 3’-AMP, pApA, and 2’3’-cAMP^29^. Moreover, SntA was proposed to work alongside other extracellular nucleotidases of *S. suis* to degrade adenine nucleotides and generate adenosine (Ado), which has been found to impact bacterial survival and virulence^30^. Lastly, a third c-di-AMP ePDE, termed CpdB, capable of degrading c-di-AMP, 2’3’-cGAMP among other nucleotides, was recently identified in *Bacillus anthracis*^31^. Loss of CpdB led to enhanced adhesion, invasion, and colonization despite a reduction in virulence in *B. anthracis* in silkworm and mouse infection models^31^.

Through BLASTp searches and structural predictions, we identified a surface-anchored extracellular phosphodiesterase (ePDE) in the *E. faecalis* reference strain OG1RF^32^ and termed it EecP for *<u>E</u>. faecalis* <u>e</u>xtracellular <u>c</u>-di-AMP <u>p</u>hosphodiesterase. Further analysis revealed that *eecP* is a part of the *E. faecalis* core genome but absent in most enterococcal species, including the other human-associated pathogen *E. faecium*. Deletion of *eecP* led to a significant accumulation of c-di-AMP in the extracellular space, confirming its role in nucleotide degradation. While inactivation of *eecP* did not alter the survival of *E. faecalis* in conditions typically associated with c-di-AMP regulation *in vitro*, the Δ*eecP* mutant strain showed increased sensitivity to killing by phagocytic cells *ex vivo*. Using two mouse infection models, we demonstrate that the impact of EecP loss and the resulting accumulation of extracellular c-di-AMP on *E. faecalis* pathogenesis may depend on the site of infection. Nevertheless, disseminated infection was more robust in animals infected with the *eecP* deletion mutant, indicating that EecP is a novel virulence factor in *E. faecalis* that cleaves extracellular c-di-AMP, likely modulating host immune surveillance.

## Results

### Identification of a cell surface-associated phosphodiesterase that is unique to *E. faecalis*

Using *in silico* analyses, we identified a putative cell wall-anchored phosphodiesterase in the genome of *Enterococcus faecalis*, designated OG1RF_RS00285, herein referred to as EecP (*<u>E.</u> faecalis* <u>e</u>xtracellular <u>c</u>-di-AMP <u>p</u>hosphodiesterase). EecP contains an N-terminal secretion signal and is predicted to be anchored to the peptidoglycan cell wall via a C-terminal LPxTG motif. Unlike other c-di-AMP-specific extracellular phosphodiesterases (ePDEs), which typically consist of a single metallophosphoesterase-nucleotidase (MP-NT) two-domain structure, EecP features duplicated MP-NT domain pairs (**Fig 1A**). Although the two MP-NT domains of EecP (MP-NT/D1 and MP-NT/D2) exhibit low amino acid identity (∼24%), they are predicted to share some structural similarity, with a root mean square deviation (RMSD) of 4.56 and a template modeling (TM) score of 0.72 (**Fig 1B-C**).

**Figure 1.**
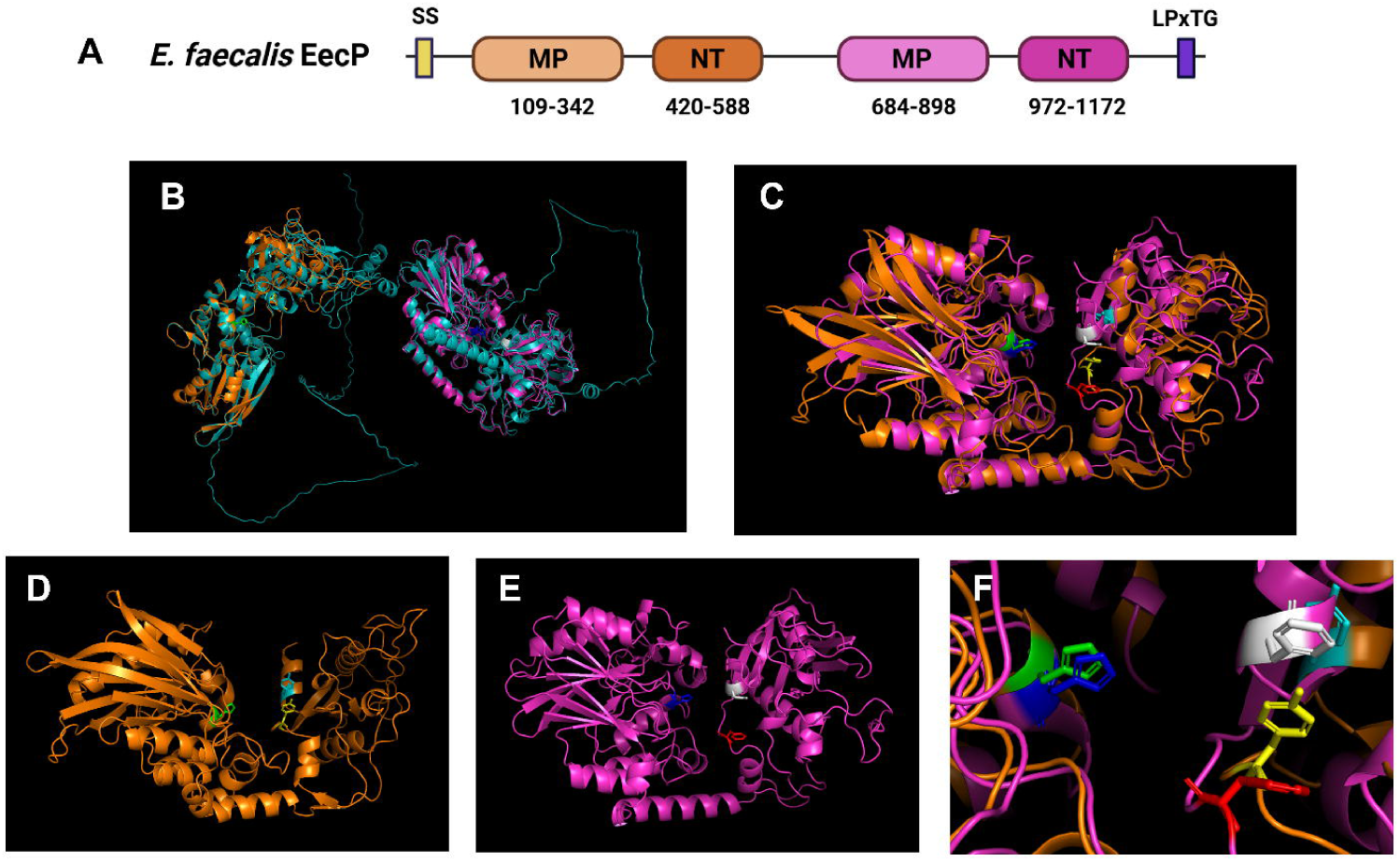
EecP is an *E. faecalis* extracellular c-di-AMP phosphodiesterase. (**A**) Domain structure of EecP, SS: secretion signal, MP: metallophosphoesterase, NT: 5’ nucleotidase, LPxTG: cell-surface anchoring. (**B**) Alpha-fold2 structure prediction of *E. faecalis* EecP (OG1RF_ RS00285) showing two distinct structures containing metallophosphoesterase (MP) and nucleotidase (NT) domain pairs (MP-NT/D1 and MP-NT/D2). MP-NT/D1 is highlighted in orange, and MP-NT/D2 in magenta. (**C**) MP-NT/D1 and MP-NT/D2 domains superimposed on each other using PyMol. (**D**) MP-NT/D1 structure highlighting the conserved histidine (H203 in green) and tyrosines (Y491 in yellow and Y580 in cyan) residues. (**E**) MP-NT/D2 structure highlighting the conserved histidine (H758 in blue) and phenylalanines (F1043 in red and F1159 in white). (**F**) Close-up of H203 (green), Y491 (yellow), and Y580 (cyan) of MP-NT/D1 and H758 (dark blue), F1043 (red), and F1159 (white) of MP-NT/D2.

The activity of bacterial nucleotidases with MP-NT domain pairs has been previously investigated in the studies of *Escherichia coli* UshA and CpdB and *S. suis* SntA ^33–35^. Collectively, these studies suggest that the MP domain (Pfam: PF00149) is responsible for enzymatic activity, while the NT domain (Pfam: PF02872) mediates substrate binding and specificity^36^. More specifically, the enzymatic activity of MP domains is attributed to a key histidyl (H) residue within a conserved ‘NHE’ motif ^20, 35–37^. Conserved ‘NHE’ motifs can be identified in the MP-NT/D1 (H203) and MP-NT/D2 (H758) domains of EecP, indicating that both MP-NT domain structures might be catalytically active (**Fig 1D-F and S1**). In addition to the ‘NHE’ motif, two tyrosine residues in DxYx**Y**xN and NN**Y**R motifs (Y530 and Y633) in the NT domain of *S. suis* SntA were modeled to bind substrates by forming a sandwich-like structure and shown experimentally to impact substrate binding efficiency and specificity^29^. While both tyrosine residues are conserved in MP-NT/D1 (Y491 and Y580), these residues are replaced by phenylalanines in MP-NT/D2 (F1043 and F1159) (**Fig 1D-F**). The presence of these key residues with similar side chain structures in EecP MP-NT/D1 and MP-NT/D2 suggests that there may be a similar function but unique specificity to these duplicated domain structures.

Because nucleotidases with MP-NT domain pair structures have been identified in both Gram-positive and Gram-negative bacteria, we performed BLASTp searches using EecP as the query sequence in NCBI and BV-BRC databases to identify homologs, with particular emphasis on cell wall-anchored proteins^38^. We find that homologs with duplicated MP-NT domain structures are present in very few closely related organisms, including *Enterococcus caccae, Enterococcus haemoperoxidus,* and *Vagococcus lutrae* (**Fig 2 and S2**). Despite the conservation of EecP across *E. faecalis* strains, no homologs were identified in *E. faecium* genomes, the second most prevalent human-associated enterococcal species. On the other hand, cell wall-anchored nucleotidases with single MP-NT domain structures can be found in other human pathogens, including *S. agalactiae*, *Streptococcus pyogenes, Staphylococcus aureus*, *Listeria monocytogenes*, and *B. anthracis* (**Fig 2 and S2**). Interestingly, the Gram-positive model organism *Bacillus subtilis* is one of the few species harboring a large surface-anchored nucleotidase, YfkN, with a duplicated MP-NT domain structure. YfkN has been shown to have phosphodiesterase activity against 2’3’-cyclic nucleotides and 5’ mononucleotides^39^.

**Figure 2.**
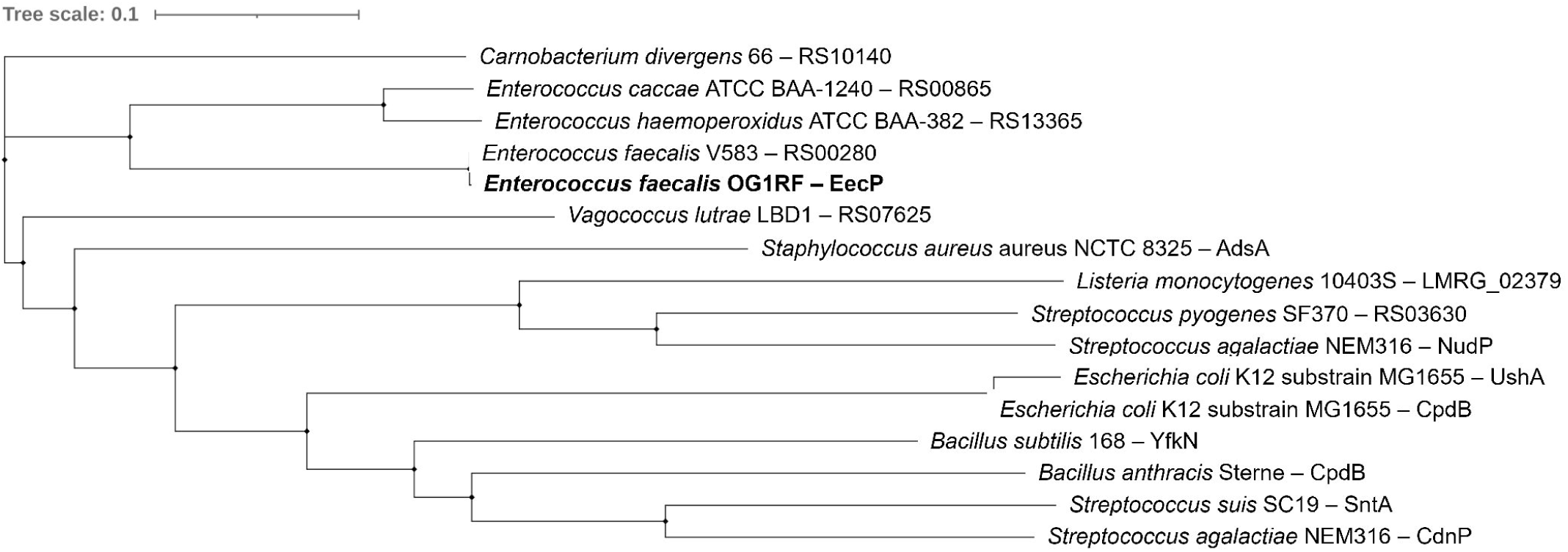
Phylogenetic analysis of EecP homologs. BLASTp searches using EecP as the query sequence in NCBI and BV-BRC were used to identify homologs in Gram-positive and Gram-negative organisms. Protein sequences were obtained from genomes of representative strains indicated for each species available in NCBI or Bio-Cyc. The phylogenetic tree was constructed using multiple sequence alignments of putative homologs using EMBL-EBI Clustal Omega and iTOL.

### *E. faecalis* is an ePDE+ organism that degrades extracellular c-di-AMP

To evaluate the ability of *E. faecalis* to degrade c-di-AMP extracellularly, we developed a thin-layer chromatography (TLC)-based assay (see methods for detail) to monitor the degradation of radiolabeled c-di-AMP using live bacterial cell suspensions. To track product formation, we used ATP and c-di-AMP standards and products formed by purified DhhP, one of the intracellular PDE in *E. faecalis*, as controls (**Fig S3A**). To assure rigor, we assessed extracellular c-di-AMP degradation in two reference *E. faecalis* strains, OG1RF and V583, that were grown to mid-log before spiking with [^32^P]-c-di-AMP in 1:10 volume ratio (1 µl c-di-AMP per 10 µl cells). We observed that both strains rapidly degraded the radiolabeled nucleotide, generating various byproducts corresponding to pApA, AMP, and inorganic phosphate (Pi) (**Fig 3**). To rule out spontaneous degradation of c-di-AMP over time, we incubated [^32^P]-c-di-AMP in the same buffer used to resuspend live cells without observing any detectable loss in the intensity of the c-di-AMP spot for at least 6 h (**Fig S3B**). These results point to the capability of *E. faecalis* cells to very rapidly degrade extracellular c-di-AMP beyond AMP at the cell surface interface. Importantly, we observed comparable results using *S. agalactiae* strains COH1 and NGBS93 (**Fig 3**), previously shown to encode the c-di-AMP ectonucleotidase CdnP^20^.

**Figure 3.**
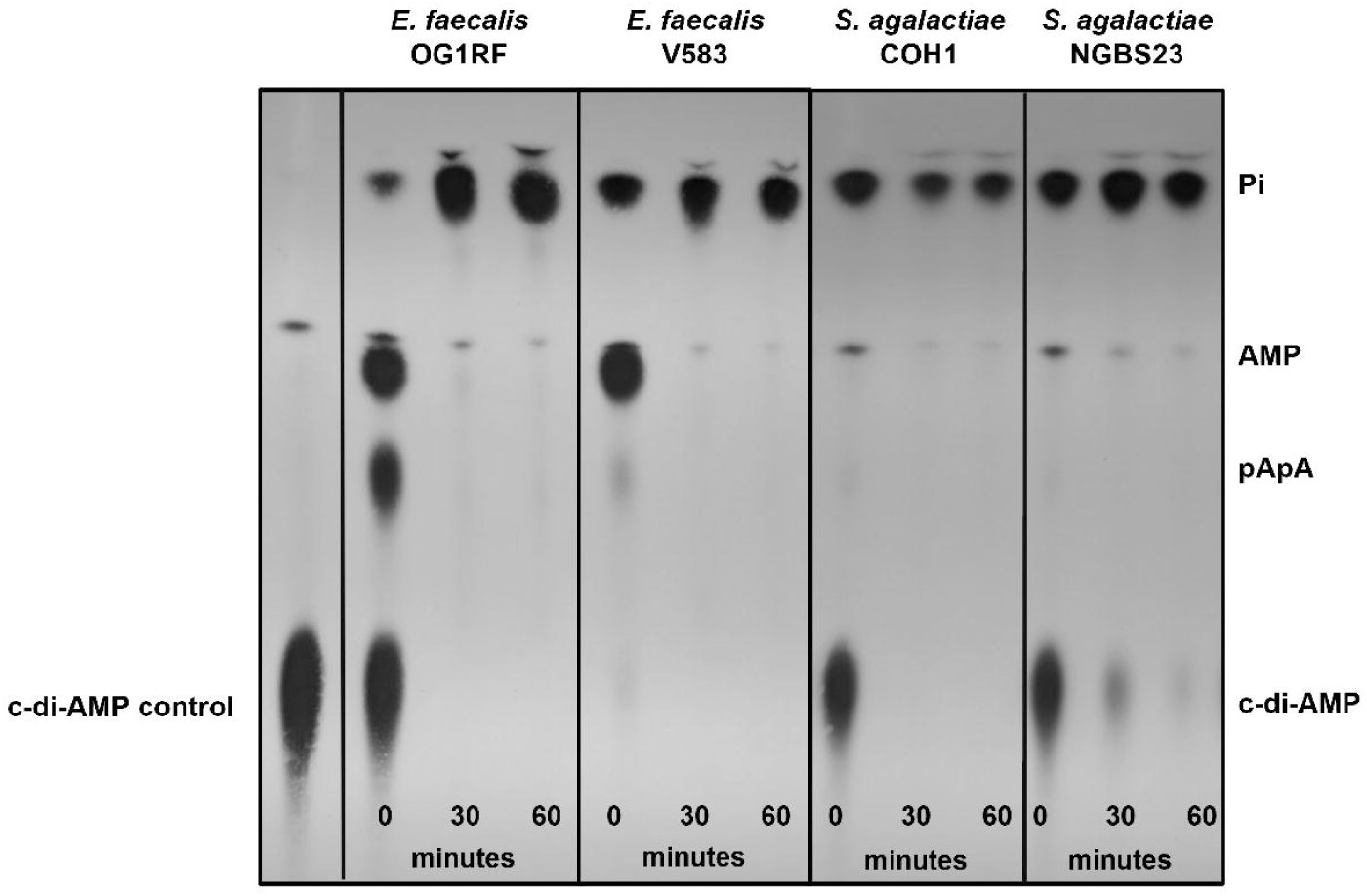
Extracellular c-di-AMP degradation in *E. faecalis* and *S. agalactiae* strains. Radiolabeled c-di-AMP was added to mid-log-grown cultures suspended in 50 mM Tris-Cl containing 5 mM MnCl_2_ and sampled over time for c-di-AMP degradation. Reaction aliquots were collected at the indicated time points and inactivated by boiling before spotting on a PEI-cellulose plate for TLC separation.

### Loss of *eecP* leads to extracellular accumulation of c-di-AMP

To investigate the possible role of EecP in c-di-AMP degradation, we used a markerless in-frame deletion strategy^40^ to generate a clean deletion strain (Δ*eecP*) in the reference strain OG1RF and the pheromone-inducible (cCF10) vector pCIE to genetically complement the mutant strain (Δ*eecP* pCIE::*eecP*)^41^. First, we used a commercially available ELISA to quantify intra- and extracellular c-di-AMP pools in the OG1RF, Δ*eecP,* and Δ*eecP* pCIE::*eecP* strains during different phases of growth^42, 43^. Briefly, cultures were grown in a chemically defined media (CDM), and both cell lysates and supernatants collected at early-log (OD_600_ 0.4), mid-log (OD_600_ 0.6), and late-log (OD_600_ 0.8) phases and processed for intra- and extracellular c-di-AMP quantifications, respectively. While there were no significant differences in intracellular c-di-AMP pools during the different growth phases among strains, extracellular c-di-AMP levels were significantly higher in Δ*eecP* supernatants when compared to OG1RF supernatants (**Fig 4A-B**). Complementation of the Δ*eecP* strain restored extracellular c-di-AMP to parent strain levels (**Fig 4C-D**). Of note, a Δ*eecP* strain harboring an empty pCIE vector was used in the genetic complementation studies to ensure that the addition of chloramphenicol and cCF10 pheromone did not interfere with the results.

**Figure 4.**
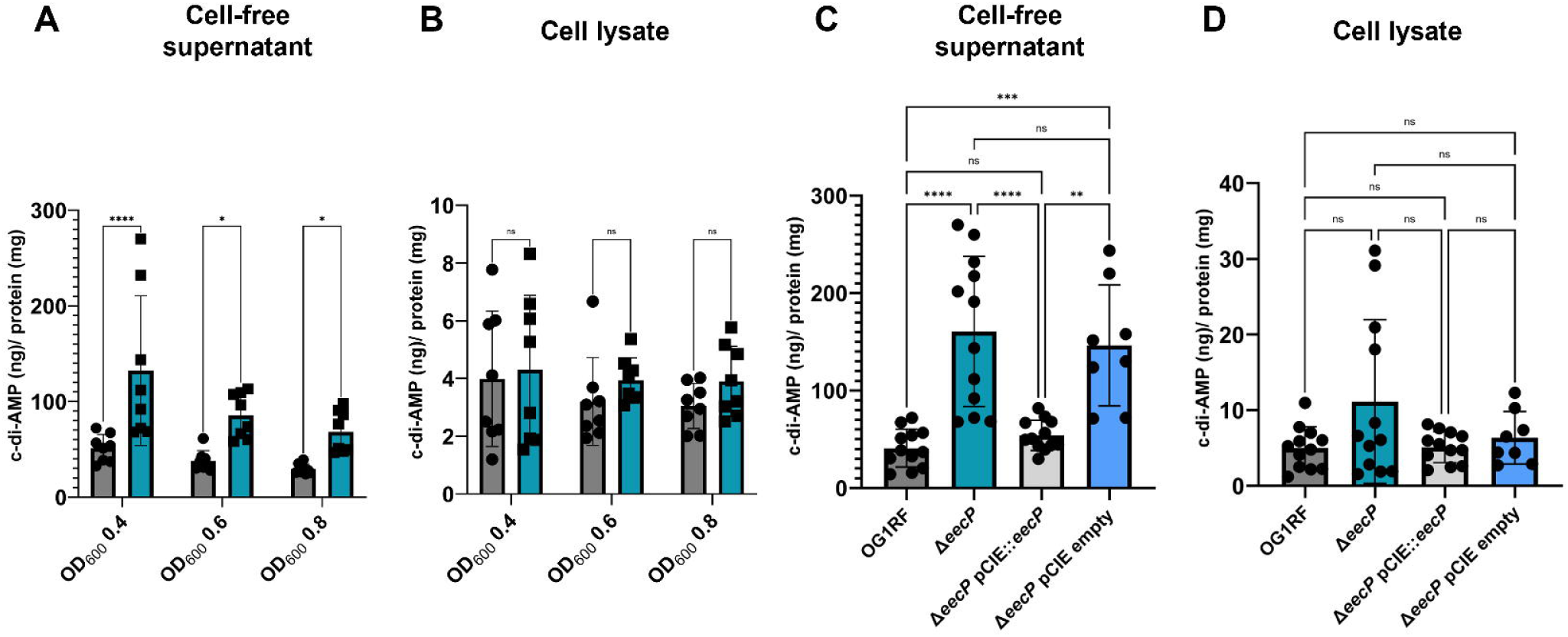
Extracellular c-di-AMP accumulation in the Δ*eecP* strain. OG1RF (parent strain) and Δ*eecP* strains were grown in CDM to OD_600_ of 0.4, 0.6, and 0.8, and c-di-AMP levels in cell-free supernatants (A) and cell lysates (B) quantified using a c-di-AMP ELISA (Cayman Chemicals). Complementation of Δ*eecP (*Δ*eecP* pCIE::*eecP* strain) restores extracellular c-di-AMP to parent strain levels in cell-free supernatants (C) and cell lysates (D). Data is derived from at least four biological replicates. Statistical analysis was performed by ordinary Two-way ANOVA, followed by Uncorrected Fisher’s LSD multiple comparisons test. ****, P≤0.0001, ***, P≤0.001, **, P≤0.01, *, P≤0.05. (ns) not significant.

To validate the ELISA results and obtain insights into the c-di-AMP degradation products of EecP, we used the TLC-based assay to monitor the degradation of [^32^P]-c-di-AMP in the presence of OG1RF and Δ*eecP* live cells. As anticipated, c-di-AMP was efficiently degraded upon contact with OG1RF cells, with no c-di-AMP detected after 30 minutes (**Fig 5**). As already shown in Figure 3, the formation of both pApA, AMP, and the release of Pi from c-di-AMP degradation can be detected immediately upon contact with OG1RF cells. While still able to degrade c-di-AMP over time, Δ*eecP* cell suspension did so much more slowly, with a considerable amount of c-di-AMP still observed after 60 minutes of incubation, and no pApA or AMP was detected on the TLC plate. Again, the complemented Δ*eecP* strain degraded radiolabeled c-di-AMP as effectively as the parent strain. Collectively, these results reveal that EecP degrades extracellular c-di-AMP, yielding pApA, AMP, and Pi.

**Figure 5.**
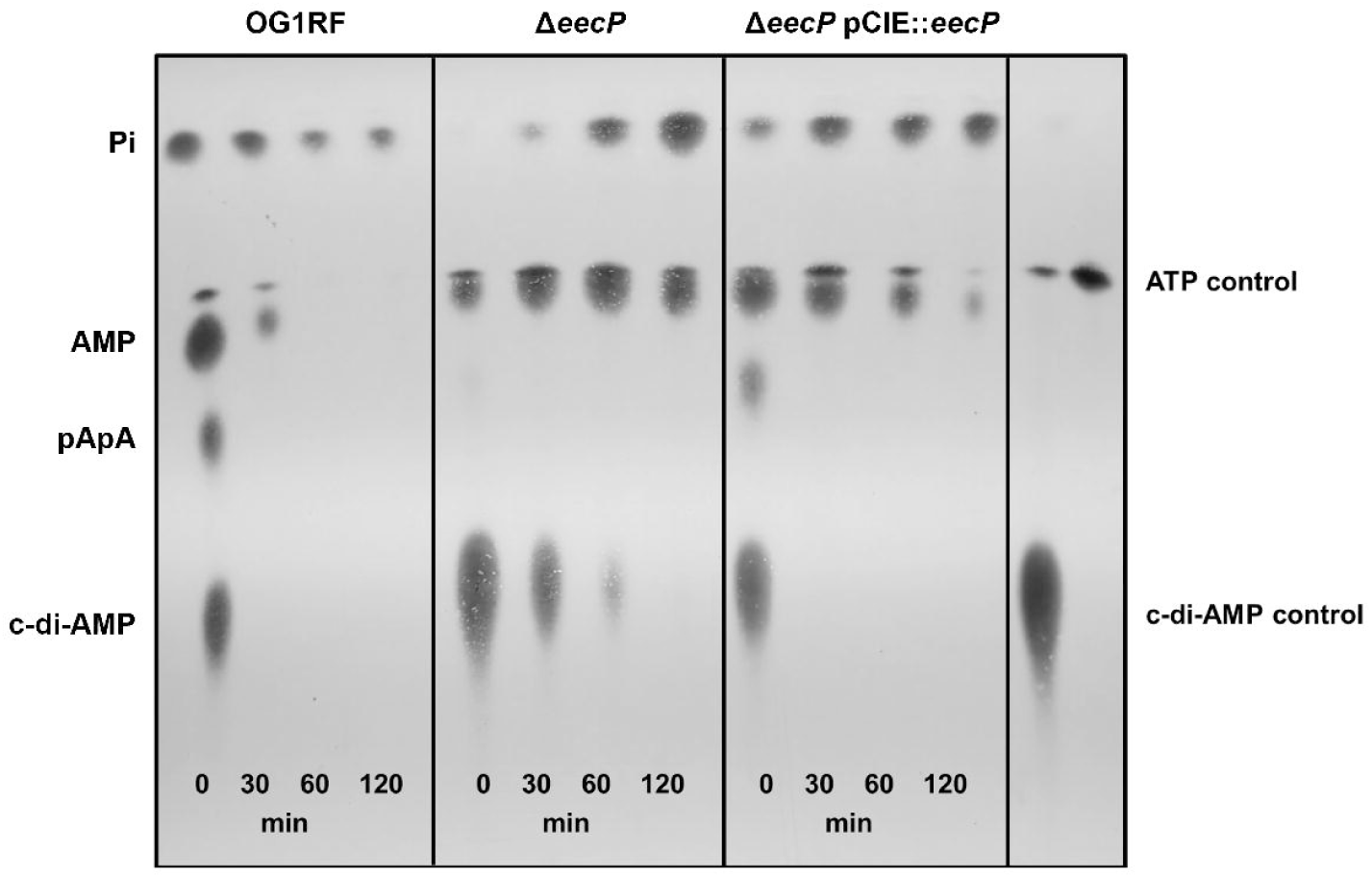
Loss of *eecP* leads to defects in extracellular c-di-AMP degradation. TLC of cell-free supernatants of OG1RF, Δ*eecP*, and Δ*eecP* pCIE::*eecP* from cultures grown to mid-log in CDM and spiked with [P]-c-di-AMP.

### Conserved histidyl residue H203 is essential for EecP activity

To determine the individual contributions of each MP-NT domain, we introduced point mutations into the *eecP* coding sequence within the pCIE::*eecP* plasmid, originally constructed for genetic complementation of the Δ*eecP* strain. Specifically, we substituted the critical histidyl residue in one or both MP-NT pairs with alanine to generate the SDM1^H203A^, SDM2^H758A^, and double mutant SDM1^H203A^SDM2^H758A^ strains (**Fig 6A**). These plasmids were then introduced into the Δ*eecP* background to assess the ability of the EecP variants to degrade c-di-AMP.

**Figure 6.**
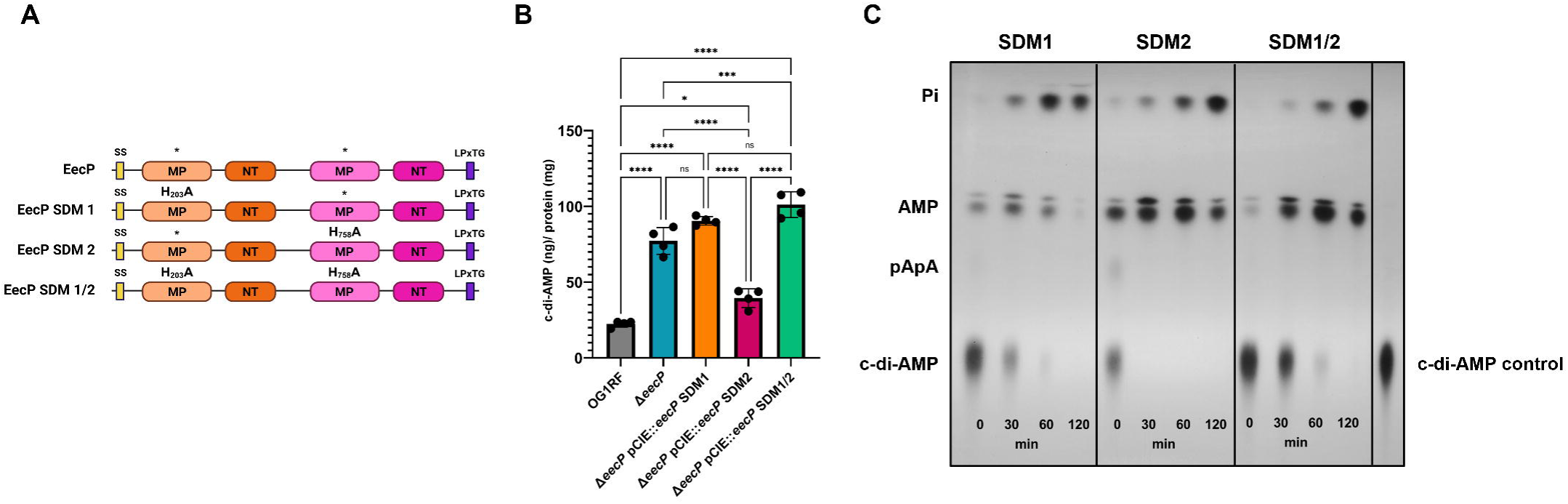
The histidine residue of the conserved NHE motif of MP-NT/D1 is essential for c-di-AMP degradation. (**A**) Domain structure of EecP and site-directed mutants (SDM) created, SS: secretion signal, MP: metallophosphoesterase, NT: 5’ nucleotidase, LPxTG: cell-surface anchoring. SDM 1 has an H203A substitution in MP-NT/D1, SDM 2 has an H758A substitution in MP-NT/D2, and SDM 1/2 has both H203A and H758A substitutions. (**B**) Strains were grown in CDM to OD_600_ 0.4 and c-di-AMP in their cell-free supernatants quantified via ELISA. Data is derived from four biological replicates. Error bars represent the standard deviation. Statistical analysis was performed by ordinary Two-way ANOVA, followed by Turkey’s multiple comparisons test. ****, P≤0.0001, ***, P≤0.001, *, P≤0.05. (ns) not significant. (**C**) TLC of cell-free supernatants of SDM 1, SDM 2, and SDM 1/2 mid-log grown cultures incubated in the presence of radiolabeled [^32^P]-c-di-AMP.

Using a quantitative c-di-AMP ELISA, we found that extracellular c-di-AMP levels in Δ*eecP* and SDM1^H203A^ strains were comparable. On the other hand, the SDM2 ^H758A^ strain exhibited only a modest increase in c-di-AMP compared to the parent strain (**Fig 6B**). Notably, extracellular c-di-AMP levels were slightly higher in the SDM1^H203A^SDM2^H758A^ strain than in SDM1^H203A^ alone (**Fig 6B**). These findings were corroborated by TLC analysis of radiolabeled c-di-AMP degradation across the different mutants (**Fig 6C**). Together, these results indicate that the first MP-NT domain is the primary contributor to c-di-AMP degradation, while the second domain plays a minor role, if any.

### The Δ*eecP* strain phenocopied the parent strain under *in vitro* conditions

Having established that EecP is responsible for extracellular c-di-AMP degradation, we next investigated the ability of the Δ*eecP* strain to grow under conditions where c-di-AMP is essential for growth or optimal proliferation. In nutrient-rich brain heart infusion (BHI) medium and chemically defined medium (CDM), the Δ*eecP* strain exhibited growth patterns very similar to that of the parent strain (**Fig 7A-B**). Similar growth between strains was observed in BHI and CDM supplemented with 0.5 M KCl, 1 mM of the osmolyte glycine betaine, or 1% peptone; all conditions previously demonstrated to require c-di-AMP for growth^15^ (**Fig 7 C-H**). Given that several cell wall-anchored proteins mediate adhesion and biofilm formation, we also assessed the ability of the Δ*eecP* strain to form biofilms *in vitro*. As seen in the growth cultures, no discernible phenotype was observed when comparing the biofilm formation of parent and Δ*eecP* strains (**Fig S4**).

**Figure 7.**
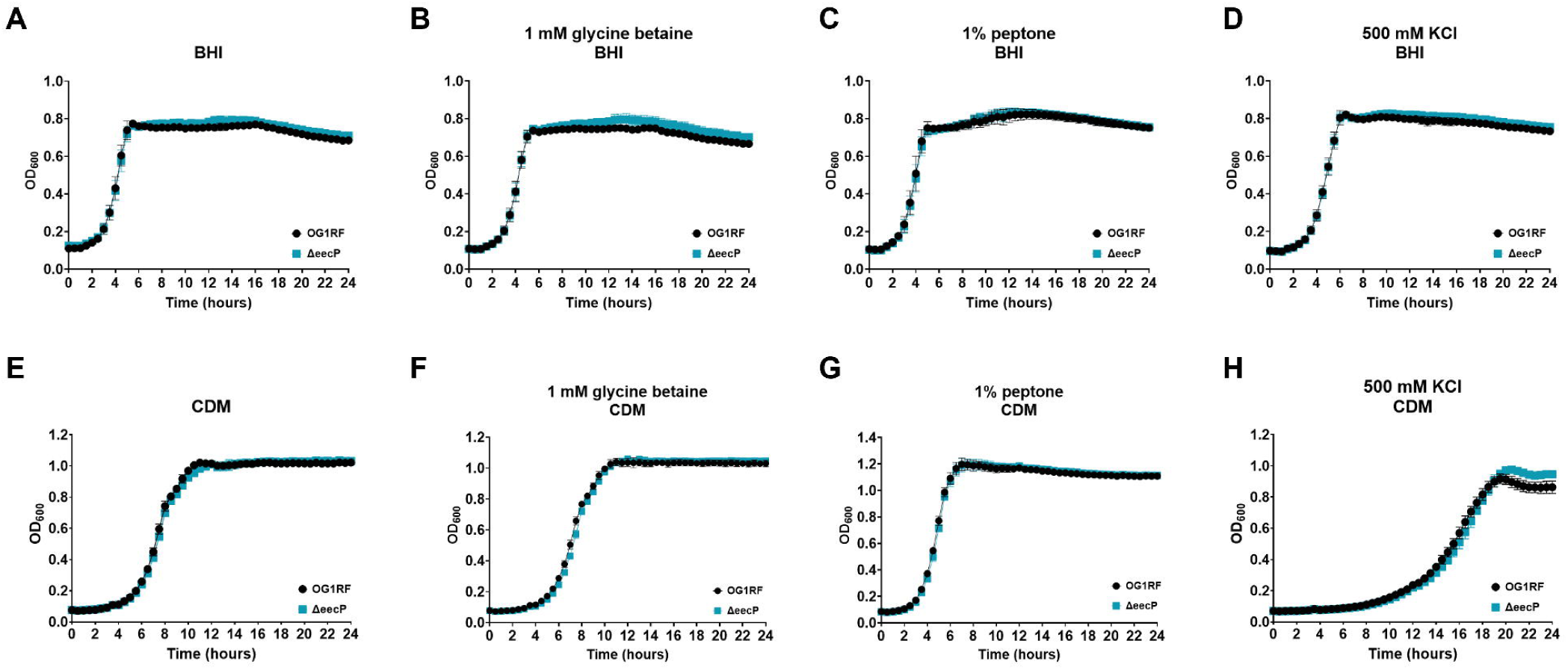
Deletion of *eecP* does not impact *E. faecalis* growth under *in vitro* conditions that depend on intracellular c-di-AMP. Growth curves of OG1RF and Δ*eecP* in brain heart infusion (BHI) (**A**) and a chemically defined media (CDM) (**E**), supplemented with 1 mM glycine betaine (**B** and **F**), 1% peptone (**C** and **G**), and 500 mM KCl (**D** and **H**). Curves represent the average derived from six biological replicates. Error bars represent standard deviation.

### Loss of *eecP* differentially impacts survival and growth in biological fluids and immune cells *ex vivo*

Although we obtained compelling evidence that EecP can degrade extracellular c-di-AMP, its role in *E. faecalis* pathophysiology remained unclear. The ability of *E. faecalis* to persist and replicate in the bloodstream and within various cell types, including macrophages, has been proposed as an important immune evasion strategy that is directly linked to its virulence^44–47^. Given that c-di-AMP is a potent pathogen-associated molecular pattern (PAMP), we hypothesized that extracellular accumulation of c-di-AMP would be relevant during host-*E. faecalis* interactions. To explore this, we first assessed the ability of the Δ*eecP* strain to grow and survive in human whole blood and serum and its susceptibility to killing by freshly isolated human neutrophils. Although deletion of *eecP* did not affect *E. faecalis* survival or growth in blood or serum *ex vivo* (**Fig 8A-B**), the survival of the Δ*eecP* strain within polymorphonuclear neutrophils (PMNs) was slightly reduced compared to the parental strain, albeit the difference was not statistically significant (**Fig 8C**). Since the *Streptococcus suis* protein SntA has been reported to interfere with complement activity by sequestering the C1q complement protein^27^, we examined whether Δ*eecP* showed altered susceptibility to rabbit complement. No differences in survival between the parental and mutant strains were observed (**Fig 8D**). Next, we assessed the ability of the Δ*eecP* strain to survive within naïve (M0) and polarized macrophages of both M1 and M2 phenotypes. Monocytes isolated from C57BL/6 mice were first differentiated into macrophages (M0) using M-CSF, followed by polarization into M1-like and M2-like macrophages using GM-CSF and IL-4, respectively. Here, the Δ*eecP* strain exhibited significantly impaired survival within M0, M1-like, and M2-like macrophages (**Fig 8E**).

**Figure 8.**
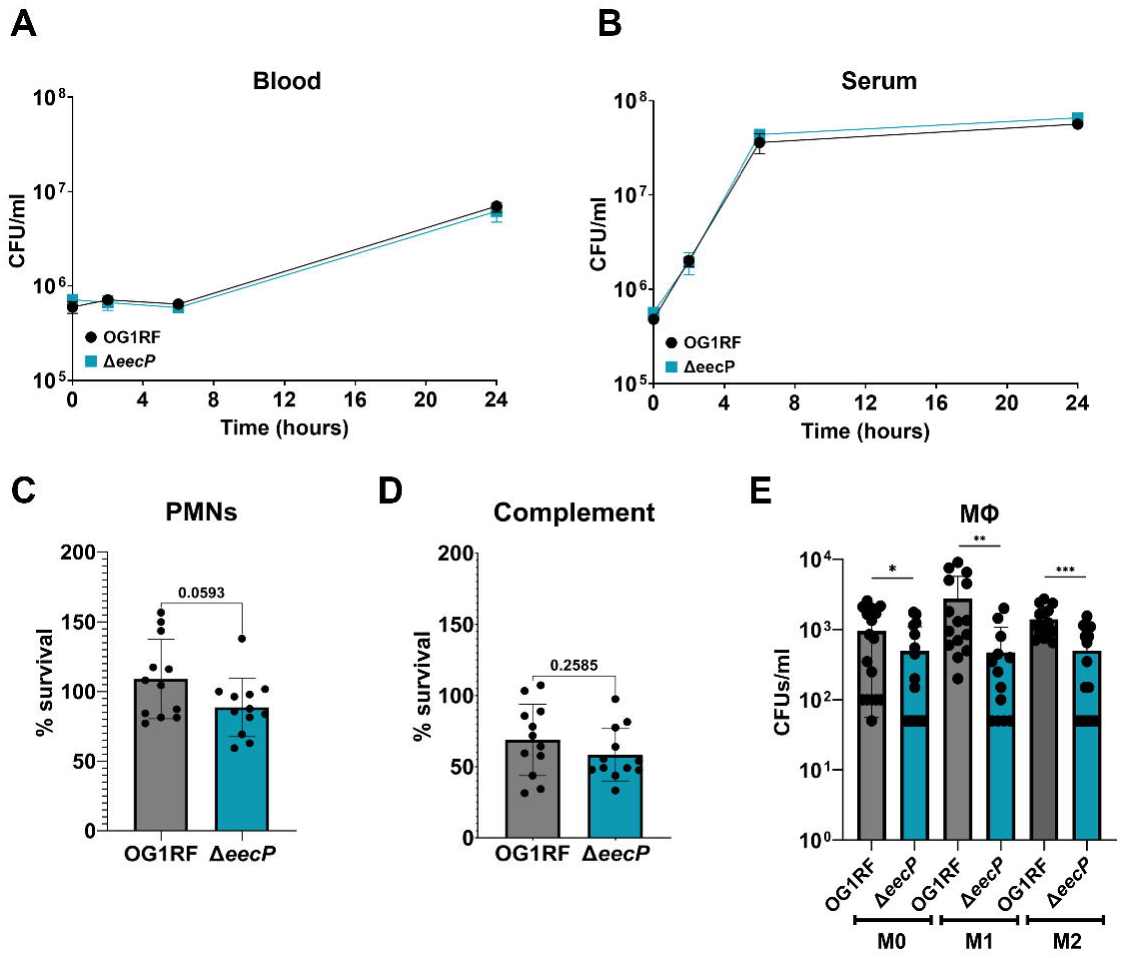
Inactivation of *eecP* differentially impacts the viability and growth of *E. faecalis ex vivo*. Growth of the parent strain OG1RF and Δ*eecP* in (**A**) human blood and (**B**) human serum. Viability of OG1RF and Δ*eecP* in (**C**) polymorphonuclear cells (PMNs), (**D**) rabbit complement, and (**E**) murine BMDMs. All experiments were conducted using at least three biological replicates. Data points shown in **C**, **D**, and **E** are the result of robust regression and outlier removal (ROUT method). Unpaired nonparametric t-test was used to determine significance, ****, P≤0.0001, ***, P≤0.001, **, P≤0.01, *, P≤0.05. Error bars represent standard deviation.

### Deletion of *eecP* impacts virulence and facilitates systemic dissemination

To directly assess the potential role of EecP in pathogenesis, we employed two established murine infection models to evaluate the virulence of the Δ*eecP* strain. First, we used a peritoneal challenge model, which leads to systemic dissemination within 24 h. No significant differences were observed in bacterial loads recovered from the peritoneal cavity of mice infected with either the OG1RF (parent) or Δ*eecP* strains at 24 or 48 h. However, higher numbers of Δ*eecP* were recovered from hearts, spleens, and livers at 24 h post-infection compared to mice infected with the parental strain (**Fig 9A-B**). After 48 h, a significant increase in bacterial burden was detected in the kidneys and spleens of Δ*eecP*-infected animals (**Fig 9B**). While not statistically significant, a trend suggesting that Δ*eecP* persists in the heart and liver in higher numbers compared to the parent strain can also be observed at 48 h post-infection (**Fig 9B**).

**Figure 9.**
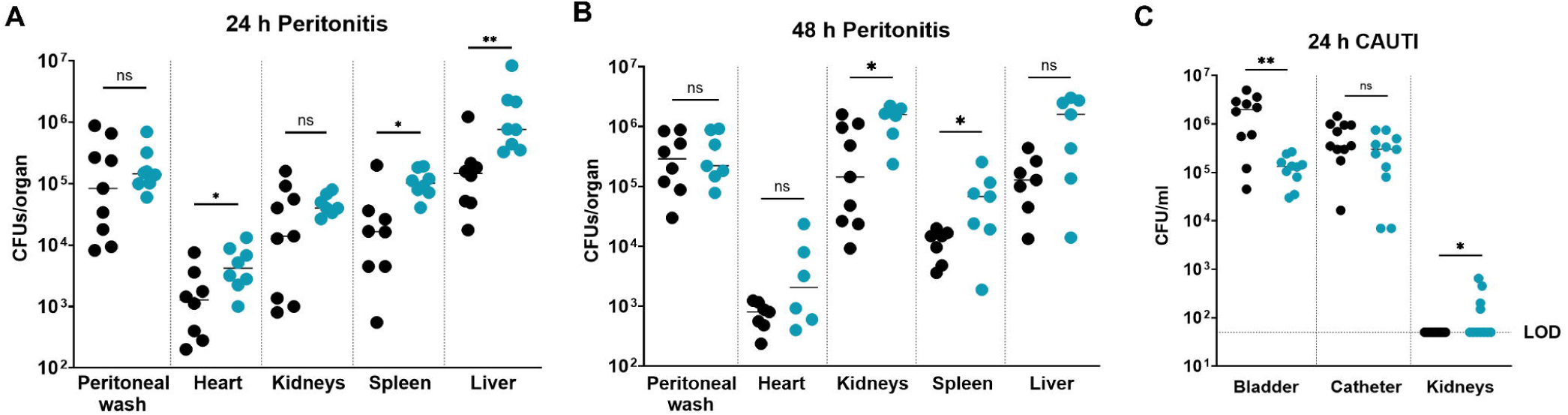
Δ*eecP* has increased dissemination in murine peritonitis and CAUTI models. (**A-B**) Recovery of *E. faecalis* OG1RF and Δ*eecP* from the peritoneal wash, heart, kidneys, spleen, and liver homogenates of mice after 24 or 48 h post-infection. (**C**) Recovery of *E. faecalis* OG1RF and Δ*eecP* from bladder, catheter implant, and kidney homogenate collected from mice 24 h post-infection. Each data point represents one animal, and the data shown is the result of robust regression and outlier removal (ROUT method). Mann Whitney unpaired nonparametric t-test was used to determine significance, **, P≤0.01, *, P≤0.05.

As we suspected that host immune responses to extracellular c-di-AMP may vary depending on the site of infection, we next utilized a catheter-associated urinary tract infection (CAUTI) mouse model^48^ to evaluate the ability of Δ*eecP* to colonize the bladder, the catheter implant, and to ascend to the kidneys. At 24 h post-infection, both OG1RF and Δ*eecP* strains were recovered in similar numbers from the implanted catheters. However, Δ*eecP* exhibited approximately a 1-log reduction in CFU recovered from the bladder (**Fig 9C**). Despite this impaired bladder colonization, the Δ*eecP* strain ascended to the kidneys in approximately 36% of infected mice, whereas no kidney colonization was observed in mice infected with the parental strain (**Fig 9C**). These findings indicate that the impact of *eecP* deletion on virulence is niche-specific. Nevertheless, the loss of *eecP* and the associated increase in extracellular c-di-AMP appear to facilitate systemic dissemination.

## Discussion

Cyclic di-AMP is a bacterial second messenger essential for the regulation of important physiological processes (e.g. osmobalace and cell wall homeostasis), stress response activation, and virulence ^9–11^. In addition, c-di-AMP is recognized by host cells as a potent PAMP; therefore, it serves to alert the host of a potential bacterial infection. In particular, c-di-AMP activates host receptors STING^20, 21, 23^, DDX41^24^, RECON^25^, and ERAdP^26^, which, in turn, activate corresponding immune signaling pathways leading to the production of inflammatory cytokines. While we have previously demonstrated that loss of intracellular c-di-AMP homeostasis drastically impairs *E. faecalis* fitness and virulence^15^, the role of extracellular c-di-AMP in *E. faecalis* interactions with the host was not investigated. In this report, we identified and characterized EecP, an ePDE unique to *E. faecalis*, that modulates extracellular c-di-AMP levels. Using *in vitro*, *ex vivo,* and *in vivo* models, we showed that while loss of *eecP* minimally affected *E. faecalis in vitro* fitness, the Δ*eecP* strain was more sensitive to killing by phagocytic cells while displaying niche-specific phenotypes in mouse infection models. Given that the enzymes that mediate c-di-AMP synthesis and degradation are considered viable therapeutic targets, a better understanding of the role and mechanisms that modulate intra- and extracellular c-di-AMP pools in bacterial pathogens such as *E. faecalis* is imperative for the rational design and development of new antimicrobial therapies.

With the identification of EecP, we place *E. faecalis* among the few bacterial pathogens known to encode c-di-AMP-specific extracellular phosphodiesterases (ePDEs). Notably, homologs of EecP are rare among other *Enterococcus* species and absent in the human-associated *E. faecium*. EecP is also unique because it contains duplicated MP-NT catalytic domains, distinguishing it from ePDEs characterized in *S. agalactiae*, *S. suis*, and *B. anthracis*. Consistent with the activity of other c-di-AMP-degrading phosphodiesterases, our TLC analysis indicates that EecP breaks down c-di-AMP into both pApA and AMP, and likely inorganic phosphate (Pi). Without EecP, *E. faecalis* appears to degrade c-di-AMP more slowly, producing AMP and, eventually, Pi. Previous studies on *S. agalactiae* revealed the presence of two ePDEs, CdnP and NudP, which act sequentially to convert c-di-AMP into AMP and subsequently into adenosine (Ado) and Pi^20, 49^. While Ado is a well-known immunosuppressive molecule linked to anti-inflammatory responses^50–52^, its accumulation in the extracellular space correlates with reduced survival and virulence in *S. agalactiae* and *S. suis* ^30, 49^. Because both pApA and AMP rapidly disappear in the presence of EecP⁺ *E. faecalis* (**Fig 5**), and EecP⁻ cells retain the ability to convert c-di-AMP into AMP and then Pi, we speculate that *E. faecalis* encodes an extracellular enzyme capable of converting AMP into adenosine with a limited activity towards c-di-AMP. Of interest, pApA formation was not observed in the presence of Δ*eecP* cells, indicating that EecP might be the only extracellular enzyme capable of generating pApA. As we have shown that *E. faecalis* can take up exogenous c-di-AMP^15^, we also cannot rule out that a fraction of radiolabeled c-di-AMP is internalized and degraded by intracellular PDEs. Future studies will be aimed at determining extracellular adenosine accumulation due to c-di-AMP degradation, and identifying additional enzymes involved in extracellular c-di-AMP and AMP metabolism.

Single amino acid substitutions of conserved histidyl residues in MP domains of EecP strongly indicate that c-di-AMP hydrolysis is primarily mediated by the MP-NT/D1. On the other hand, a conserved motif of the substrate-binding NT domain in MP-NT/D1 contains two tyrosines, while these tyrosine residues are replaced by two phenylalanine residues in MP-NT/D2. Of note, tyrosine and phenylalanine are aromatic amino acids that only differ by the presence of a hydroxyl group on the aromatic side chain of tyrosine. Notably, the tyrosine residues of EecP MP-NT/D1 are conserved in CpdB-like PDEs, including *E. coli* CpdB, *S. agalactiae* CdnP, *B. anthracis* CpdB, and *S. suis* SntA (**Fig S1**) that, despite some differences in substrate and cofactor specificity, have been shown to degrade c-di-AMP ^20, 29, 31, 53–56^. Alternatively, the phenylalanine residues of EecP MP-NT/D2 are conserved in *S. agalactiae* NudP and *E. coli* UshA, which have been shown to degrade 5’ mononucleotides^49, 57^ (**Fig. S1**). These subtle differences in key residues of NT domains in MP-NT/D1 and MP-NT/D2 hint that the duplicated MP-NT domains may have different substrate specificities. While lacking direct evidence, we speculate that EecP MP-NT/D2 might be functionally related to the *S. agalactiae* NudP and can hydrolyze AMP into adenosine.

As previously observed with the *S. agalactiae* Δ*cdnP* strain^20^, the inactivation of *eecP* did not yield noticeable phenotypes *in vitro*, indicating that extracellular accumulation of c-di-AMP in single cultures of *E. faecalis* has no significant effects. To explore the biological significance of EecP, we then tested the ability of the mutant to survive within phagocytic cells and cause disease using two mouse infection models. Survival assays using PMNs, PMNs, and murine BMDMs indicated that the Δ*eecP* strain is more susceptible to phagocytic killing, albeit the differences observed were only significant within BMDMs. In *S. agalactiae*, the viability of the *cdnP* mutant within THP-1 cells or BMDMs was not altered when compared to the parent strain, despite evidence that the Δ*cdnP* strain induced a stronger IFN-β response^20^. In the intraperitoneal challenge model, we observed increased bacterial burden in distal organs of animals infected with the Δ*eecP* strain but not in the peritoneal wash retrieved from the initial site of infection. In the CAUTI model, the Δ*eecP* strain colonized the bladder more poorly when compared to the parent strain. Still, despite this apparent attenuation in virulence, the infection ascended to the kidneys only in mice infected with Δ*eecP*. These results indicate that while the impact of the loss of EecP on disease outcome might depend on where the infection is initiated, extracellular c-di-AMP accumulation facilitates systemic dissemination.

As it is already known, sensing of c-di-AMP by host receptors can induce various immune signaling pathways that will culminate in the production of a myriad of cytokines like IFN-β, IL-1β IL-6, and TNFα^58^. While the induction of these cytokines is intended to limit bacterial proliferation and contain infection, their activation can be detrimental and increase host susceptibility to infections. For example, several studies have reported that type I interferons (IFNs), known to be induced by c-di-AMP, enhance the susceptibility of mice to *Listeria monocytogenes* infection via impaired bacterial clearance, a reduction in proinflammatory immune cell populations, upregulation of apoptotic genes, and enhanced T cell death ^59–63^. Also induced by c-di-AMP, IL-6 is a cytokine with a central role in Th17 cell differentiation, cells associated with protection against extracellular pathogens^64, 65^. Despite its protective role, IL-6 induction has been deemed an immune evasion strategy by *Mycobacterium tuberculosis* by inhibiting autophagy, a key cellular homeostasis mechanism involved in innate defense against bacterial and viral infections^66^. Increased IL-6 has also been proposed to favor *E. faecalis* intracellular survival in macrophages^67^. RECON, a cytosolic pattern recognition receptor (PRR), relieves repression of NF-κB upon binding c-di-AMP, promoting the transcription of genes involved in innate and adaptive immunity^25^. Interestingly, *E. faecalis* has been shown to suppress NF-κB activation in macrophages, a phenomenon linked to promoting polymicrobial catheter-associated urinary tract infection (CAUTI) pathogenesis^68^. Together, these examples illustrate how excess c-di-AMP and activation of downstream responses can enable bacterial persistence and increase host susceptibility.

The dysregulation of host immune responses due to the pressure of sustained c-di-AMP may help explain the enhanced systemic dissemination of the Δ*eecP* strain. Furthermore, the examples described above underscore the complexity and nuanced interplay of immune responses to c-di-AMP. While the targeted identification of host factors that sense and respond to c-di-AMP has yielded important insights, the application of global “omics” approaches will be invaluable for deepening our understanding of c-di-AMP as a bacterial pathogen-associated molecular pattern (PAMP) molecule and its broader role in host-pathogen interactions.

In this study, we provide the first in-depth characterization of the *E. faecalis* extracellular phosphodiesterase, termed EecP, a novel enzyme that degrades c-di-AMP extracellularly. In addition, EecP emerged as an important virulence factor impacting infection dissemination in two murine models. These findings underline the importance of extracellular c-di-AMP in host-pathogen interactions and highlight the role of this molecule as an active immunomodulator during *E. faecalis* infection.

---

**Figure S1.** Conservation of key residues among MP-NT-containing enzymes. Sequence alignment of MP-NT containing enzymes in select Gram-positive and Gram-negative bacteria. Alignment was performed with Multiple Sequence Comparison by Log-Expectation (MUSCLE) via SnapGene. Conserved residues highlighted (in blue) for EecP MP-NT/D1 and MP-NT/D2 include H203 and H758 (**A**), Y491 and F1043 (**B**), and Y580 and F1159 (**C**).

**Figure S2.** Domain structures of EecP homologs. BLASTp searches using EecP as the query sequence in NCBI and BV-BRC were used to identify homologs in Gram-positive and Gram-negative bacteria. Protein sequences were obtained from genomes of representative strains indicated for each species available in NCBI or Bio-Cyc. The phylogenetic tree was constructed using multiple sequence alignments of putative homologs using EMBL-EBI Clustal Omega and iTOL.

**Figure S3.** Recombinant *E. faecalis* DhhP (rDhhP) degrades c-di-AMP into AMP. (**A**) 50 µM of purified rDhhP was mixed with radiolabeled [^32^P]-c-di-AMP in 50 mM Tris-HCl 5 mM MnCl_2_ buffer. Reaction aliquots were collected at the indicated time points and inactivated by boiling before being resolved via TLC. (**B**) [P]- c-di-AMP suspended in reaction buffer was incubated at 37° for 30 min before inactivation by boiling. The inactivated suspension was then incubated at room temperature and spotted to check for spontaneous degradation for up to 6 h.

**Figure S4.** Biofilm biomass quantification of parent strain OG1RF and Δ*eecP* grown in 96-well plates in BHI for 24 h. Data points represent fifteen biological replicates from five independent experiments.

**Table S1.** Chemically defined media formulation

**Table S2.** Primers used in this study.

## Materials and methods

### Ethics statement

Animal procedures for murine infections were approved by the University of Florida Institutional Animal Care and Use Committee (protocol # 202200000241) and the University of Notre Dame Institutional Animal Care and Use Committee (protocol # 22-01-6971). All animal care was consistent with the Guide for the Care and Use of Laboratory Animals from the National Research Council and the USDA Animal Care Resource Guide.

### Bacterial strains and growth conditions

The bacterial strains used in this study are listed in **Table 1**. *E. faecalis* strains were grown in brain heart infusion (BHI) or chemically defined media (CDM) at 37°C under static conditions. The CDM formulation is outlined in **Table S1** in the supplementary material. Strains bearing the pCIE plasmids were grown under antibiotic pressure (10 μg/ml chloramphenicol) with 5 ng/ml of cCF10 pheromone (Mimotopes) added for gene activation. To determine growth kinetics, overnight cultures grown in BHI or CDM were adjusted to an optical density at 600 nm (OD_600_) of ∼0.2-0.3 and inoculated into fresh media at a ratio of 1:100. Growth was monitored at an OD_600_ using an automated growth reader for up to 24 h. Additional components KCl (Fisher Scientific), glycine betaine (Sigma Aldrich), and peptone (Thermo Scientific) were added to BHI or CDM media to concentrations indicated in the figures.

**Table 1.**
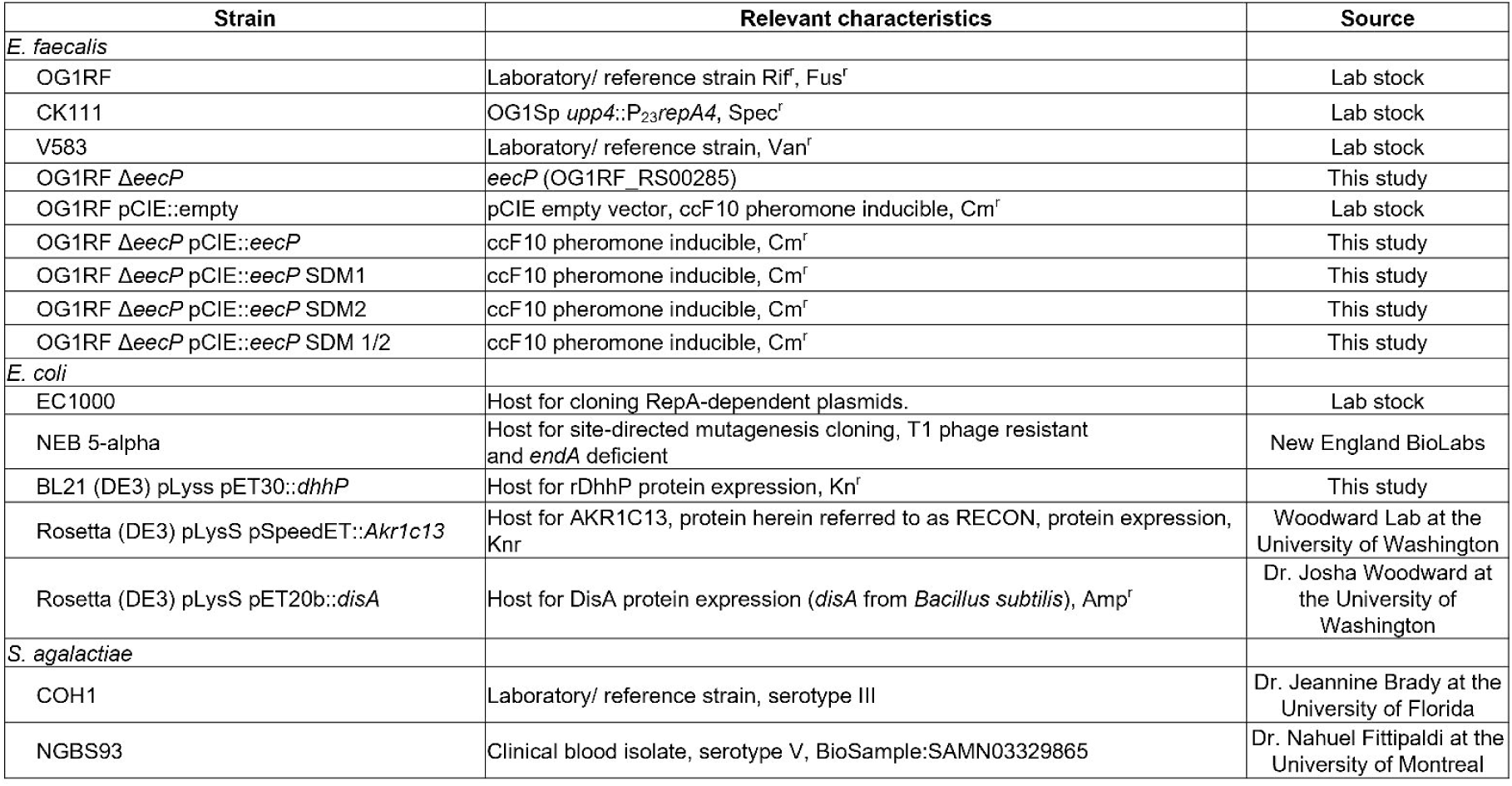
Strains used in this study.

| Strain | Relevant characteristics | Source |
| --- | --- | --- |
| *E. faecalis* |  |  |
| OG1RF | Laboratory/ reference strain Rif<sup>r</sup>, Fus<sup>r</sup> | Lab stock |
| CK111 | OG1Sp *upp4::P<sub>23</sub>repA4*, Spec<sup>r</sup> | Lab stock |
| V583 | Laboratory/ reference strain, Van<sup>r</sup> | Lab stock |
| OG1RF $\Delta eecP$ | *eecP* (OG1RF_RS00285) | This study |
| OG1RF pCIE::empty | pCIE empty vector, ccF10 pheromone inducible, Cm<sup>r</sup> | Lab stock |
| OG1RF $\Delta eecP$ pCIE::*eecP* | ccF10 pheromone inducible, Cm<sup>r</sup> | This study |
| OG1RF $\Delta eecP$ pCIE::*eecP* SDM1 | ccF10 pheromone inducible, Cm<sup>r</sup> | This study |
| OG1RF $\Delta eecP$ pCIE::*eecP* SDM2 | ccF10 pheromone inducible, Cm<sup>r</sup> | This study |
| OG1RF $\Delta eecP$ pCIE::*eecP* SDM 1/2 | ccF10 pheromone inducible, Cm<sup>r</sup> | This study |
| *E. coli* |  |  |
| EC1000 | Host for cloning RepA-dependent plasmids. | Lab stock |
| NEB 5-alpha | Host for site-directed mutagenesis cloning, T1 phage resistant and *endA* deficient | New England BioLabs |
| BL21 (DE3) pLyss pET30::*dhhP* | Host for rDhhP protein expression, Kn<sup>r</sup> | This study |
| Rosetta (DE3) pLysS pSpeedET::*Akr1c13* | Host for AKR1C13, protein herein referred to as RECON, protein expression, Knr | Woodward Lab at the University of Washington |
| Rosetta (DE3) pLysS pET20b::*disA* | Host for DisA protein expression (*disA* from *Bacillus subtilis*), Amp<sup>r</sup> | Dr. Josha Woodward at the University of Washington |
| *S. agalactiae* |  |  |
| COH1 | Laboratory/ reference strain, serotype III | Dr. Jeannine Brady at the University of Florida |
| NGBS93 | Clinical blood isolate, serotype V, BioSample:SAMN03329865 | Dr. Nahuel Fittipaldi at the University of Montreal |

### General cloning techniques

The nucleotide sequence of *eecP* (OG1RF_RS00285) was obtained from the *E. faecalis* OG1RF genome via NCBI. The Wizard Genomic DNA Purification kit (Promega) was used for isolation of bacterial genomic DNA (gDNA), and the Monarch Plasmid Miniprep kit (New England Biolabs) was used for plasmid purification. The Zymo DNA Clean and Concentrator (Zymo Research) was used to isolate PCR products. Colony PCR was performed using ProMega GoTaq Green PCR 2X master mix (Promega) with primers listed in **Table S2**.

### Construction of gene deletions and genetically complemented strains

Deletion of *eecP* in *E. faecalis* OG1RF was carried out using the pCJK47 markerless genetic exchange system^40^, as previously described. Briefly, approximately 1 kb size sequences upstream and downstream of *eecP* were amplified using the primers listed in **Table S2**. The two amplicons were ligated to the pCJK47 vector, linearized with XbaI and EcoRI, and transformed into *E. coli* EC1000. Then, the vector containing the ligated amplicons was electroporated into *E. faecalis* CK111 (donor strain) that was used to deliver the plasmid to the recipient *E. faecalis* OG1RF strain, where markerless genetic exchange occurred. The gene deletion was confirmed by PCR screening the gene site and flanking sequences, followed by Sanger sequencing and whole-genome sequencing to ensure no additional mutations emerged. For plasmid complementation, the *eecP* gene was amplified by PCR using the primers listed in **Table S2** and ligated into the pCIE vector linearized with BamHI and HindIII restriction enzymes to yield the plasmid pCIE::*eecP*. The plasmid was propagated in *E. coli* EC1000, verified by sequencing, and electroporated into the *E. faecalis* Δ*eecP* strain as described elsewhere^40^.

### Construction of site-directed mutant complemented strains

The vector pCIE::*eecP* was isolated from the propagating *E. coli* EC1000 strain. Then, using the Q5 Site-Directed Mutagenesis Kit (New England Biolabs), a histidine to alanine substitution was performed following the manufacturer’s instructions. Briefly, the substitution was created by incorporating the nucleotide change in the center of the forward primer, including at least 10 complementary nucleotides on the 3’ side of the mutation, and ensuring the 5’ ends of the two primers annealed back-to-back when designing the reverse primer. Following PCR amplification of the plasmid containing the substitution, the reaction was incubated with an enzyme mix containing a kinase, a ligase, and DpnI. These enzymes ensure the circularization of the PCR product and the removal of the template DNA. The resulting plasmid was transformed into NEB 5-alpha competent *E. coli* (New England Biolabs) and, after isolation of the plasmid from the propagating strain, electroporated into *E. faecalis* OG1RF Δ*eecP*. The mutagenized plasmids were confirmed by Sanger sequencing and whole plasmid sequencing.

### Protein expression and purification

Recombinant 6xHis-tagged *E. faecalis* DhhP, *B. subtilis* DisA, and mRECON proteins were expressed and purified from strains listed in **Table 1**. Briefly, bacterial cultures transformed with expression plasmids were grown in Luria Broth (LB) containing 0.04% (w/v) glucose and 300 µg/ml kanamycin for 15-18 h and sub-cultured to an OD_600_ of 0.05 and allowed to grow past OD_600_ 0.5 before induction. Protein expression was induced using the allolactose homolog isopropyl ß- D-1-thiogalactopyranoside (IPTG) at a concentration of 500 µM for 3 h at 30°C. Induction was confirmed by SDS-PAGE, followed by Coomassie blue staining. After expression, cell pellets were collected by centrifugation and stored at −20°C. Pellets were resuspended in equilibration buffer (20 mM NaH_2_PO_4_, 300 mM NaCl, 10 mM imidazole pH 7.5) and added protease inhibitor cocktail (Halt Protease Inhibitor Cocktail from ThermoScientific). Protein lysates were obtained by sonication using a 10-minute cycle of alternating pulses at 10% amplitude. Soluble protein fractions were collected by centrifugation and used to purify the recombinant protein by affinity chromatography using a nickel-nitrilotriacetic acid (Ni-NTA) matrix. Briefly, clear lysate was combined with Ni-NTA agarose (Marvelgent Biosciences, US) and allowed to rotate at 4°C for 2 h. The agarose-lysate mix was then added to a crystal column and allowed to settle by gravity. After collecting flow-through lysate, the column was washed thrice with three different buffer preparations: wash buffer 1: 20 mM NaH_2_PO4, 300 mM NaCl, 25 mM imidazole, 1% Triton x-100, wash buffer 2: 20 mM NaH_2_PO4, 500 mM NaCl, 25 mM imidazole, and wash buffer 3: 20 mM NaH_2_PO4, 300 mM NaCl, 50 mM imidazole. Protein was eluted four times with 200 mM NaH_2_PO_4_, 300 mM NaCl, and 250 mM imidazole. Protein elution fractions were concentrated using the Pierce Protein Concentrator PES 10K MWCO (5-20 ml) (ThermoScientific), followed by desalting/ buffer exchange using the Zeba spin columns 7K MWCO (ThermoScientific). The buffer used for the exchange for DhhP was 50 mM Tris-Cl, 5 mM MnCl_2_. Concentrated DisA and mRECON were buffer exchanged with DisA activity assay buffer containing 40 mM Tris pH 7.5, 0.1 M NaCl, 20 mM MgCl_2_. A final concentration step was performed using the Pierce Protein Concentrator PES 10K MWCO (100-500 µl) (ThermoScientific).

### Protein structure modeling

Domain mapping and annotation were performed with Pfam, now InterPro. AlphaFold Server was used to generate all prediction structures. PyMol was used for structure alignment and highlighting of structures and residues. RMSD and TM scores were determined using the Protein Data Bank (PDB) Pairwise Structure Alignment tool.

### c-di-AMP quantification by ELISA

Overnight cultures in CDM were diluted in fresh media and allowed to reach the desired OD_600_. To quantify extracellular c-di-AMP, cultures were pelleted by centrifugation, and supernatant was collected. Supernatants were centrifuged again and then filtered using 0.2 μm filters (Cytiva). For intracellular c-di-AMP quantification, pellets were collected by centrifugation, washed thrice with PBS, and resuspended in 50 mM Tris-HCl pH 7.5. Cell suspensions were transferred to screwcap tubes containing glass beads and lysed by bead beating, followed by heat inactivation at 95°C for 5-10 min. Then, samples were centrifuged, and supernatants were collected for intracellular c-di-AMP quantification. A commercial c-di-AMP ELISA (Cayman Chemicals) was used for nucleotide quantification following the manufacturer’s instructions. Intra- and extracellular samples were diluted as needed to interpolate c-di-AMP concentration in the standard curve.

### Synthesis of [α^-32^P] labeled c-di-AMP

Radiolabeled c-di-AMP was synthesized as follows: 1 μM [α^-32^P]-ATP (Revvity, Massachusetts, US) was incubated with 1 μM of the diadenylate cyclase protein DisA in 40 mM Tris pH 7.5, 100 mM NaCl, 20 mM MgCl_2_ binding buffer at 37 °C overnight. The mixture was boiled for 5 min, followed by centrifugation to remove precipitated DisA. Synthesized [^32^P]-c-di-AMP was further purified from the mixture using recombinant His-tagged mRECON, a c-di-AMP binding protein, by affinity purification. Then, 50 μM mRECON was bound to Ni-NTA resin (Marvelgent Biosciences) for 30 min on ice with agitation. The resin was washed twice with the binding buffer to remove unbound mRECON. The resulting resin was incubated with the crude [^32^P]-c-di-AMP for 30 min at room temperature, followed by 10 min on ice. After removing the supernatant, the Ni-NTA resin was washed twice with ice-cold binding buffer and incubated with 250 μl of binding buffer for 10 min at 95 °C. The slurry was then transferred to a mini spin column (ThermoScientific, US) and centrifuged for 1 min to elute [^32^P]-c-di-AMP.

### c-di-AMP degradation assay and thin layer chromatography (TLC)

Overnight cultures in CDM were diluted in fresh media and allowed to reach mid-log (OD_600_ 0.4-0.5). Pellets were collected by centrifugation, washed thrice with 50 mM Tris-HCl pH 7.5, 5 mM MnCl_2_, and resuspended to a cell density of 10^8^-10^9^ CFU/ml. To the cell suspension, [^32^P]-c-di-AMP was added in a 1:10 volume ratio (1 µl c-di-AMP per 10 µl cells). Cell suspensions were incubated at 37°C, aliquots collected at selected time points, and immediately centrifuged to collect supernatants, followed by heat inactivation for 5 min at 95°C. For TLC, polyethylenimine (PEI) cellulose F (EMD Millipore) plates were soaked in 90% methanol for 20 min and then air dried. Then, 3-5 µl of reaction products were spotted on PEI cellulose plates and separated using a buffer containing 1:1.5 (v/v) saturated (NH_4_)_2_SO_4_ and 1.5 M NaH_2_PO_4_ pH 3.4 as a mobile phase. Air-dried TLC plates were exposed to X-ray films, and radioactive spots were visualized using a film processor.

### Growth and survival in human blood and serum

Blood was purchased from LifeSouth Community Blood Center (Gainesville, FL). This procedure was approved and performed in compliance with the University of Florida Institutional Review Board (IRB) Study #IRB 202100899. The experiments were performed with pooled blood and isolated serum from two donors of the same blood type. For isolation of pooled sera, pooled whole blood was left to stand at room temperature for 30 min, and centrifuged at 2000 rpm for 10 min. The resulting serum supernatant was aliquoted and stored at −20°C for future use. For blood and serum infections, overnight cultures grown in BHI were pelleted by centrifugation, washed thrice in PBS, normalized to an OD_600_ of 0.5, and inoculated at a 1:1,000 ratio into blood or serum. Samples were incubated at 37°C with constant rotation for blood and serum samples. At specific time points, samples were serially diluted in PBS and plated on BHI agar to enumerate colony-forming units (CFU).

### Isolation of neutrophils (PMNs)

Human neutrophils were isolated as previously described^69^. Buffy coats from 6 to 8 donors were purchased from LifeSouth Community Blood Center and processed on the day of the experiment. Briefly, an equal volume of dextran-heparin buffer (3% (wt/v) Dextran-500, 0.00568% (wt/v) of heparin, 0.9% NaCl) was added to each donor’s buffy coat and incubated at 37°C for 1 h. After incubation, the high leukocyte and low erythrocyte supernatant was transferred to a new tube. Then, the leukocyte pellet of each donor was collected by centrifugation at 500 g for 10 min at 4°C and pooled together in 30 ml of dextran-heparin buffer. The pooled leukocyte suspension was underlaid with 10 ml of Histopaque 1077 (density, 1.077 g/ml) and centrifuged at 400 g for 40 min at room temperature to isolate the granulocytes and any remaining erythrocytes (PMN-erythrocyte layer). The supernatant above this layer was discarded, and the PMN-erythrocyte cell pellet was resuspended in 1% (wt/vol) ammonium chloride for 2 to 3 min for hypotonic lysis of remaining erythrocytes. The complete lysis of erythrocytes and neutrophil isolation was achieved by repeating the hypotonic lysis step and centrifugation at 400 g for 5 min at 20°C. Trypan blue staining was used to determine the number of live neutrophils.

### PMNs infection and complement survival assay

Approximately 1 x 10^6^ freshly isolated human polymorphonuclear cells (PMNs) were resuspended in Roswell Park Memorial Institute (RPMI) 1640 medium (ThermoScientific) supplemented with 15% (v/v) inactivated fetal bovine serum (FBS) (Gibco) and added to a 24-well flat-bottom tissue-culture treated plate. PMNs were infected with ∼ 1 x 10^7^ *E. faecalis* cells for a multiplicity of infection (MOI) of 1:10 neutrophil to bacteria ratio. For complement survival studies, ∼ 1 x 10^7^ cells were inoculated in RPMI supplemented with 15% FBS and 10% rabbit complement (Rockland Immunochemicals). After the addition of bacteria, plates were incubated in a shaking incubator at 37°C with agitation at 50 rpm for 90 min. After incubation, the percentage of bacterial survival was calculated by enumerating the CFU of bacteria in the wells that have neutrophils or complement compared to bacteria-only controls.

### Isolation and polarization of bone-marrow-derived monocytes (BMDMs)

Bone marrow from femurs and tibias of 6- to 8-week-old female C57BL/6 mice was flushed with Roswell Park Memorial Institute (RPMI) 1640 media supplemented with 1X L-glutamine, 5.95 g/L HEPES, and 1 mM sodium pyruvate (VWR) and filtered using a 40 µm pore size cell strainer (VWR). Peripheral blood mononuclear cells (PBMCs) were separated from red blood cells and neutrophils using centrifugation separation with density gradients Histopaque-1077 (Sigma-Aldrich). After centrifugation, the PBMCs layer was isolated and washed with 1X DPBS. Then, PBMCs were resuspended in Dulbecco’s Modified Eagle Medium (DMEM) supplemented with 4.5 g/L glucose, 2 mM L-glutamine, 1 mM sodium pyruvate (VWR), 10% FBS, 100 units/m/ penicillin and 100 µg/ml streptomycin and incubated in cell culture flasks (Greiner Bio-One) for 24 h at 37°C, 5% CO_2_. After 24 h, 25 ng/ml recombinant mouse M-CSF (R&D Systems) was added to the culture for 5-7 days for differentiation into macrophages. Uninduced (M0) BMDMs were cultured in their respective media without cytokine treatment. For M1 polarization, cells were treated for 24 h with 100 ng/ml GM-CSF (Shenandoah Biotechnology), and with 100 ng/ml IL-4 (PeproTech) for M2 polarization.

### Antibiotic protection assay BMDMs

Approximately 2 x 10^4^ BMDMs were seeded per well in a 96-well flat-bottom tissue-culture treated plate (Greiner Bio-One). Macrophages were stimulated with the listed concentrations as described above. During the 24 h stimulation, macrophages were cultured in DMEM supplemented with 10% FBS and 1X penicillin/streptomycin and incubated at 37°C, 5% CO_2_. After 24 h, the media was removed, and the cells were washed thrice with 1X DPBS and replaced with fresh media free of antibiotics. Cells were then infected with *E. faecalis* strains at an MOI ratio of 1 and incubated for 45 min at 37°C in a 5% CO_2_ incubator to allow bacteria uptake. After 45 min, supernatants were transferred into a 96-well round bottom plate for serial dilution and CFU determination. Macrophages were washed thrice with 1X DPBS before being replaced with DMEM supplemented with 200 µg/ml gentamicin and 1X penicillin/streptomycin. After incubation at 37°C, 5% CO_2_ for 3 h, media was removed, and the cells were washed three times with 1X DPBS before treatment with distilled water for 15 min to lyse macrophages. Water was then transferred into a 96-well plate for dilutions, plated on selective BHI media containing rifampicin and fusidic acid, and incubated overnight at 37°C for CFU determination.

### Intra-peritoneal challenge mouse model

Murine peritonitis experiments were performed under protocol #202200000241, approved by the University of Florida Institutional Animal Care and Use Committee (IACUC). The murine peritonitis infection model has been described elsewhere^70^. Briefly, bacterial strains for inoculums were grown in BHI to an OD_600_ 0.5. Pellets were collected by centrifugation, washed thrice in PBS, and suspended at ∼2 × 10^8^ CFU/ml. Seven-week-old C57BL6/J mice purchased from Jackson laboratories were intraperitoneally injected with 1 ml of bacterial suspension and euthanized by CO_2_ asphyxiation 24 or 48 h post-infection. To collect the peritoneal wash, the abdomen was opened to expose the peritoneal lining, and 5 ml of cold PBS were injected into the peritoneal cavity, with 4 ml retrieved as the peritoneal wash. Following peritoneal wash collection, the spleen, kidneys, liver, and heart were surgically removed, briefly washed in 70% ethanol, and rinsed in sterile PBS. Spleens, kidneys, and hearts were homogenized in 1 ml of PBS and livers in 2 ml of PBS. All sample homogenates were serially diluted in PBS and plated on selective BHI agar plates containing rifampicin and fusidic acid.

### Murine catheter-associated urinary tract infection model

Mice were subjected to transurethral implantation of a silicone catheter and inoculated as previously described^48^. Briefly, ∼6-week-old female C57BL/6J mice purchased from the Jackson Laboratory were anesthetized by isoflurane inhalation and implanted with a 6-mm-long silicone catheter (Braintree Scientific). Immediately following catheter implantation, animals were infected with 50 μl of ∼1 × 10^6^ CFU/ml of bacterial suspension in PBS. Bacteria were introduced into the bladder lumen by transurethral inoculation. Mice were euthanized 24 h post catheterizations and infections by anesthesia inhalation, followed by cervical dislocation. Catheters were collected, and the bladder, kidneys, spleen, and heart were aseptically harvested. The organs were homogenized, and catheters were cut into small pieces and sonicated for CFU enumerations. All homogenates were plated on selective BHI agar plates containing fusidic acid and rifampicin.

## Acknowledgments

We thank Dr. Joshua Woodward at the University of Washington for the *E. coli* expression strains, Dr. Jeannine Brady at the University of Florida for the COH1 *S. agalactiae* strain, and Dr. Nahuel Fittipaldi at the University of Montreal for the NGBS93 *S. agalactiae* strain. This study was supported by NIH-NIAID R01 AI172179 to J.A.L. D.N.B. was supported by NIH-NIDCR Training Grant T90 DE021990.

## Notes

### Competing Interest Statement

The authors have declared no competing interest.

