## Supplementary figures and images for "Characterization of a Novel Cell Wall-Associated Nucleotidase of *Enterococcus faecalis* that Degrades Extracellular c-di-AMP"

### Supplemental Figure 1

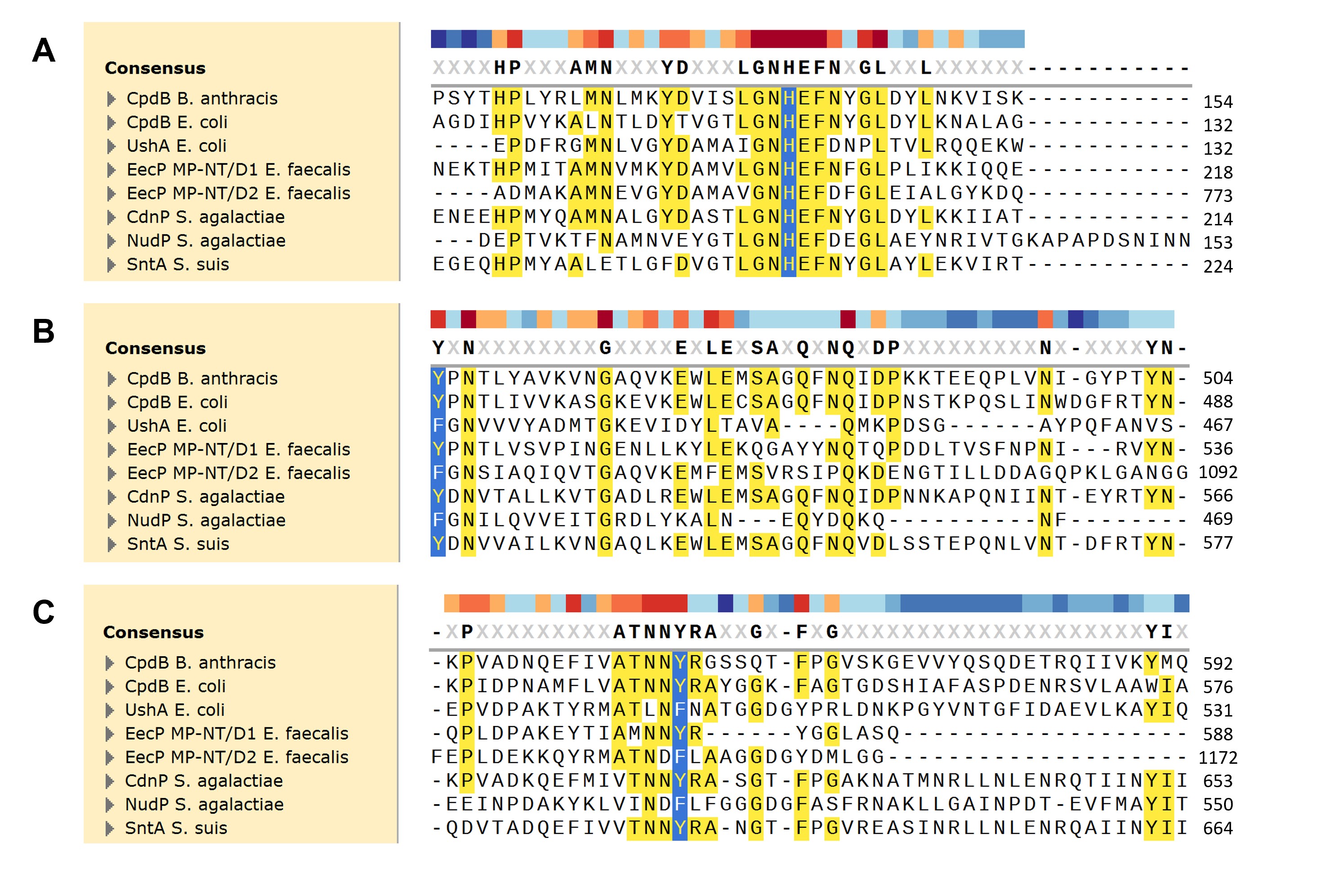

### Supplemental Figure 2

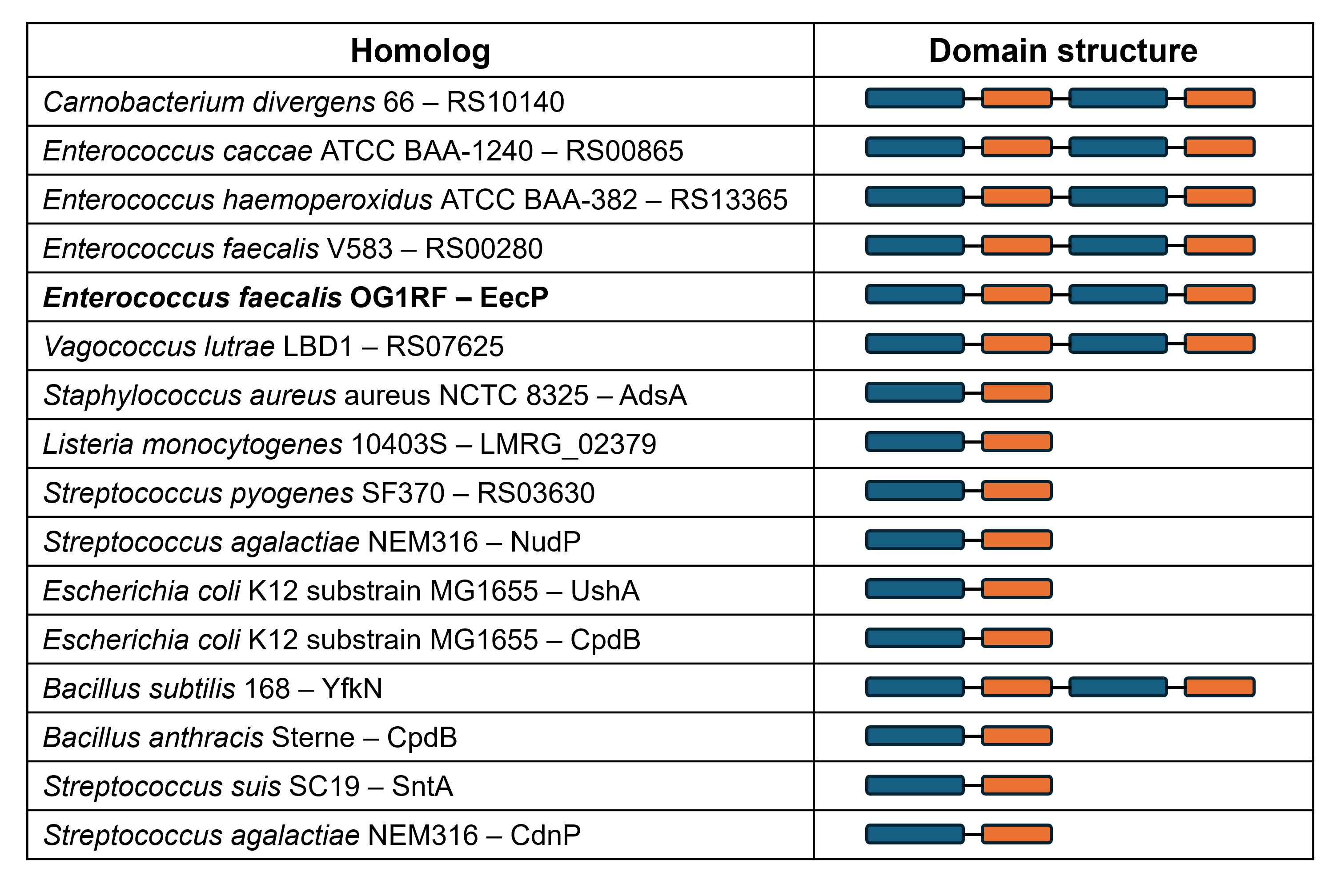

### Supplemental figure 3

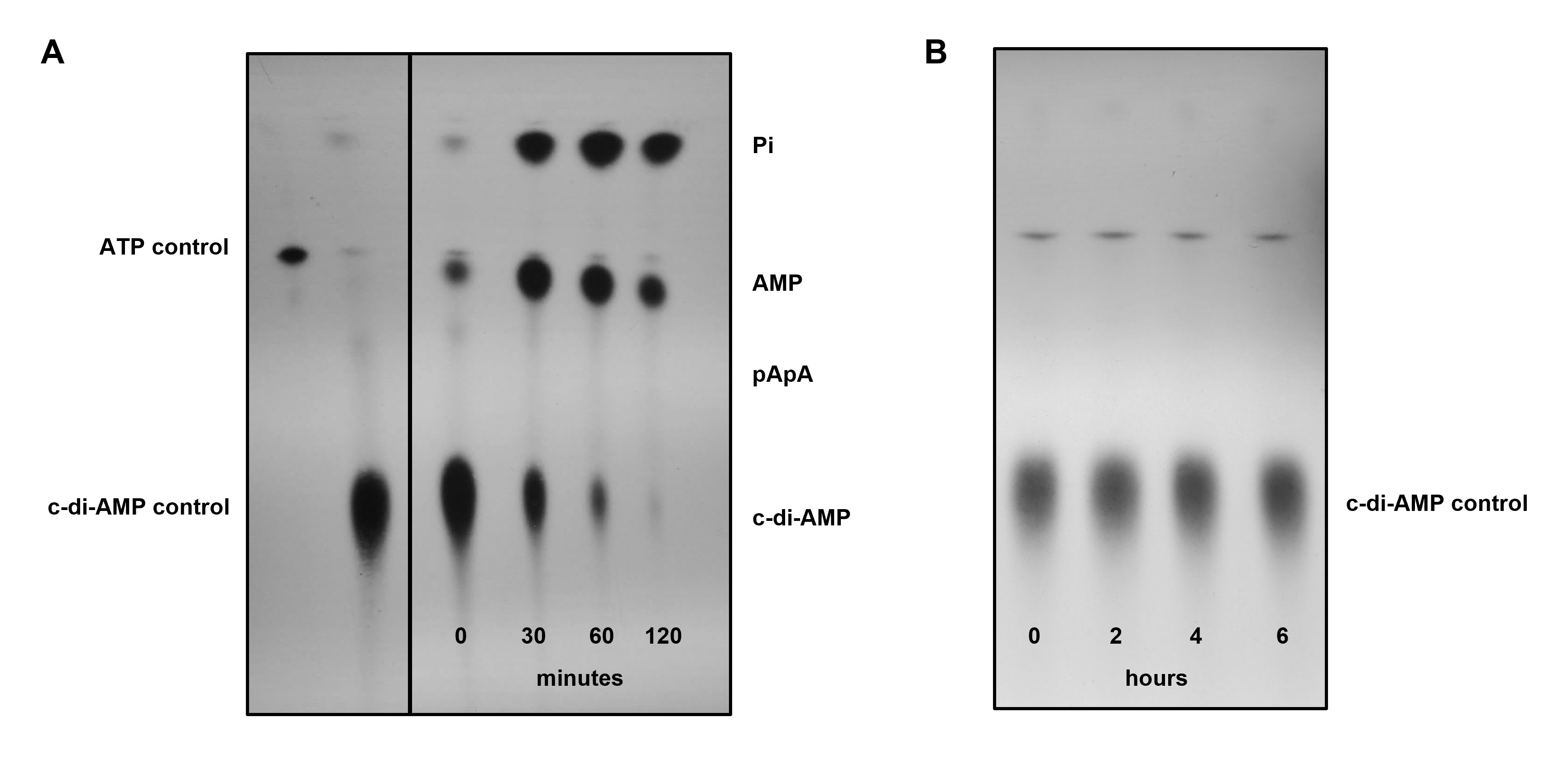

### Supplemental figure 4

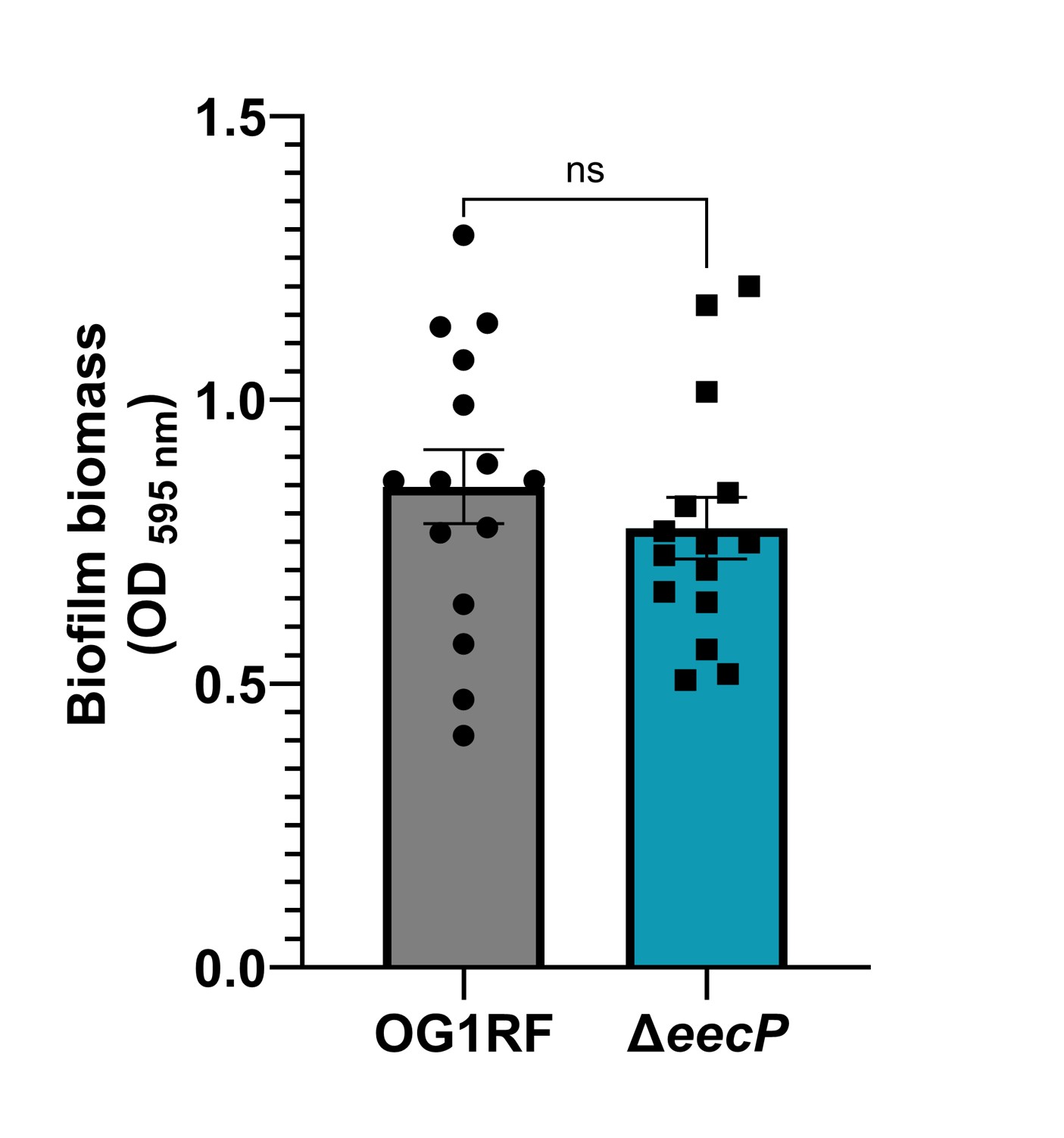
